# Inference of the core gene regulatory network underlying seam cell development in *Caenorhabditis elegans*

**DOI:** 10.1101/2023.11.28.569021

**Authors:** Alicja Brożek, Arianna Ceccarelli, Andreas Christ Sølvsten Jørgensen, Mark Hintze, Michalis Barkoulas, Vahid Shahrezaei

**Affiliations:** Department of Life Sciences, Imperial College, London SW7 2AZ, UK; Department of Mathematics, Imperial College London, Exhibition Rd, South Kensington, London SW7 2BX, UK; Mathematical Institute, University of Oxford, Oxford OX2 6GG, UK; I-X Centre for AI In Science, Imperial College London, White City Campus, 84 Wood Lane, London W12 0BZ, UK

## Abstract

Gene regulatory networks are fundamental in cellular decision-making, yet even in well-studied systems, their topologies are often poorly characterised. The nematode *Caenorhabditis elegans* contains a population of stem-like cells, known as seam cells. While seam cells are essential to generate the majority of the animal epidermis as well as specific neurons, the architecture of the underlying gene network has not been elucidated. Here, we combine experiments, mathematical modelling and statistical inference to uncover the architecture of the seam cell gene network focusing on three core transcription factors (TFs), the GATA factors ELT-1, EGL-18 and the Engrailed homolog CEH-16. We use single-molecule FISH (smFISH) to quantify TF mRNA abundance in single seam cells in both wild type and mutant backgrounds. We then predict potential TF interactions and their quantitative strengths using a combination of Modular Response Analysis, ordinary differential equations and a Bayesian model discovery approach. Taken together, our findings suggest new relationships between core TFs in seam cells and highlight an approach that can be used to infer quantitative networks from smFISH data.

## Introduction

Describing the mechanisms of stem cell patterning is fundamental in achieving an accurate understanding of tissue homeostasis. The model nematode *Caenorhabditis elegans* is well known for its robust development with high consistency in cell numbers and types of cells produced. A key cell population in *C. elegans* that exhibits stem cell-like behaviour and generates the majority of its epidermis are the so-called seam cells. Over the four larval stages of *C. elegans* post-embryonic development, the seam cells divide at regular points. The vast majority of their divisions are asymmetric and produce one seam cell and one hypodermal cell or neuroblast (Figure 1A)(Sulston and Horvitz, 1977). *C. elegans* hatches with 10 cells per lateral side, but during the L2 larval division (Figure 1A), several seam cells also divide symmetrically to increase their population from 10 to 16 (Joshi et al., 2010; Sulston and Horvitz, 1977). It is largely unknown how seam cells make robust choices between undergoing self-renewal or differentiating and how this relates to cell lineage choices and developmental time points (Hintze et al., 2020; Katsanos et al., 2017). Given the stem cell behaviour and experimental tractability of the system, the epidermis of *C. elegans* provides an ideal invertebrate model to dissect the mechanisms underlying stem cell regulation and developmental robustness.

**Figure 1.**
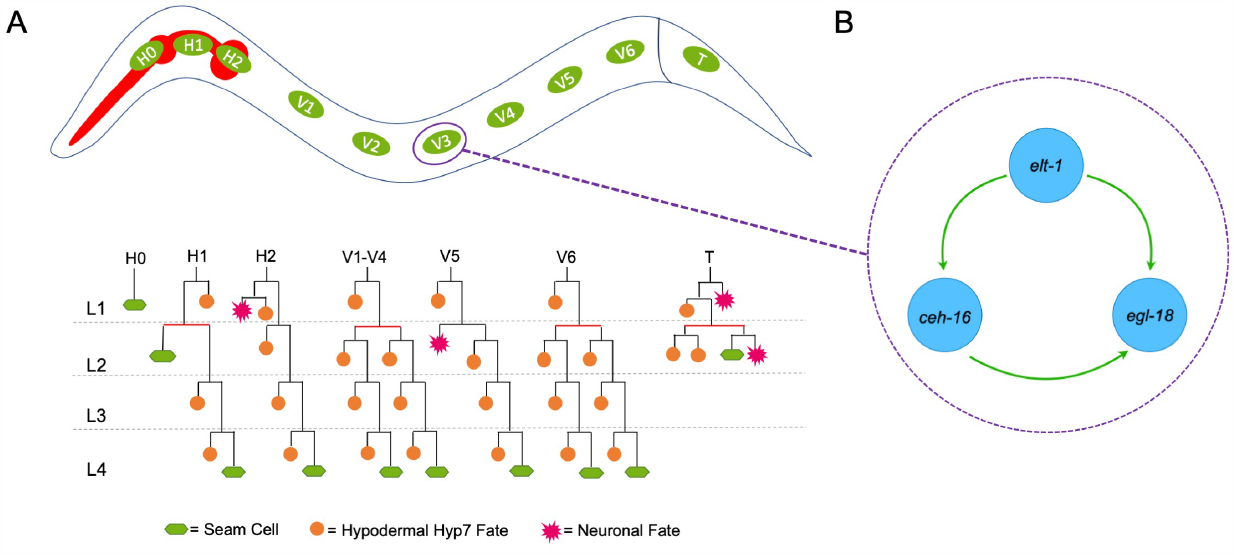
Introduction to seam cells and the core TF seam cell gene network. **A:** An illustration of the stereotypic division patterns of seam cells. Seam cell fate is denoted in green, hypodermal cell fate is denoted in blue, and neuronal fate is shown in magenta. Red bars indicate symmetric divisions. **B:** A schematic of the gene network compiled from interactions proposed in previous literature: ELT-1 activates both *ceh-16* and *egl-18* expression, and CEH-16 activates *egl-18* expression.

Genetic analysis has revealed a number of genes that are involved in the regulation of seam cell behaviour. Amongst these, three core transcription factors (TFs) are known to be essential for seam cell development, namely *elt-1, ceh-16* and *egl-18* (Gorrepati et al., 2013; Huang et al., 2009; Smith et al., 2005). ELT-1 is a GATA factor whose function is linked to seam cell fate specification and maintenance. Complete knockouts of *elt-1* are unable to specify an epidermis and thus are lethal. *elt-1* is expressed in the whole epidermis in the embryo and then localises to the seam cells post-embryonically (Spieth et al., 1991; Baugh et al., 2005; Cohen et al., 2015). This protein is known to regulate other seam-cell-specifying TFs like BRO-1, which is part of the BRO-1/RNT-1 (CBFβ/RUNX) complex that influences the choice of cells to divide symmetrically (Brabin et al., 2011).

CEH-16 is a homolog of the Drosophila gene *engrailed* as well as the human En1 and En2 (Genestine et al., 2015). The complete knockout of *ceh-16* is embryonic lethal. The gene is expressed in the AB lineage and plays a role in embryonic seam cell development (Cassata et al., 2005). During post-embryonic development, *ceh-16* is specifically expressed in the seam cells where it is required for the L2 symmetric seam cell division (Huang et al., 2009). In turn, *ceh-16* overexpression results in seam cell hyperplasia through the conversion of normally asymmetric divisions to symmetric ones (Huang et al., 2009).

EGL-18 is another GATA factor that is thought to act downstream of the Wnt signalling pathway (Koh and Rothman, 2001; Gorrepati et al., 2013). In asymmetric seam cell divisions resulting in the anterior cell acquiring a hypodermal fate, EGL-18 is thought to promote seam cell fate in posterior seam cells partly by repressing the differentiation programme, a function which is shared with CEH-16 (Cassata et al., 2005; Gorrepati et al., 2013). Strong *egl-18* loss-of-function mutations are not lethal, however, this may be due to the compensation by the paralogue GATA factor ELT-6 (Koh et al., 2002). Nevertheless, in absence of *egl-18*, a number of cells lose their ability to maintain the seam cell fate (Gorrepati et al., 2013; Koneru et al., 2021).

While there is some evidence to suggest that *elt-1, egl-18* and *ceh-16* may interact (Cassata et al., 2005; Thompson et al., 2016), the exact architecture of their interaction network has so far not been elucidated. The literature portrays ELT-1 as an early specification factor promoting the expression of both *ceh-16* and *egl-18* (Thompson et al., 2016). It additionally describes CEH-16 as a promoter of *egl-18*. This is summarised in (Figure 1B) and in Thompson et al. (2016). Understanding the way these core TFs interact and their reciprocal regulation is essential for understanding the mechanisms underlying the patterning rules for this stem cell population.

Inference methods for gene regulatory networks (GRNs) are dictated by the type of data that is available and the type of network abstraction or GRN model that one wishes to assemble (Klipp et al., 2005; Gardner and Faith, 2005; Karlebach and Shamir, 2008; Hecker et al., 2009; Mercatelli et al., 2020; Stumpf, 2021). The data used could include protein-DNA, protein-protein interaction data or gene expression data at the bulk or single-cell level. One of the simplest levels of abstraction for a GRN model is a static graph, capturing gene-gene regulatory or correlational links. An advantage of static models is that they contain fewer parameters; thus, their inference scales well computationally to larger networks. In order to tell apart correlation from causal relationships, perturbation-based data such as gene expression in genetic mutants could be used. One method to develop a directed graph model for a GRN using such data, with a quantitative measure of interaction strength, is the Modular Response Analysis (MRA) (Kholodenko et al., 2002). MRA allows for the construction of a quantitative static network using expression data from perturbation experiments without requiring detailed mechanistic information on the chemical reactions involved.

In contrast to static models, dynamic GRN models can include mechanistic information and are more useful in producing predictions. These include logical or Boolean models, ordinary differential equation (ODE) models and stochastic discrete models simulated using the Gillespie algorithm (Gardner and Faith, 2005; Karlebach and Shamir, 2008; Klipp et al., 2005). Dynamic GRNs typically have many unknown parameters and thus are harder to infer, particularly for larger networks. The task is to infer both the architecture and the parameters of such GRN models from available data. This kind of inference problem is sometimes called equation or model discovery (Brunton et al., 2016; Alahmadi et al., 2020; Both et al., 2021). A novel Bayesian model discovery approach called SLING (Sparse Likelihood-free Inference using Gibbs) uses sparsity-inducing priors to find the simplest model among a library of related models that can explain a given set of data (Jørgensen et al., 2023). This approach, which is simulation-based and related to Approximate Bayesian Computation (ABC) (Beaumont, 2019), is applicable to non-linear or stochastic models of biochemical networks. The method has been used for the inference of GRNs based on synthetic data (Jørgensen et al., 2023), but it remains to be applied to real data.

In this study, we use single-cell snapshot transcription data combined with mathematical and statistical analysis to systematically infer a core TF GRN for the seam cells in *C. elegans*. To understand how disruption in the activity of each core TF might affect the expression pattern of other genes within the network, we use single-molecule Fluorescent In Situ Hybridisation (smFISH) (Raj et al., 2008), which provides information on mRNA expression at the single cell level in wild type and loss-of-function mutants. Next, we use a modified version of the MRA algorithm to test links between components in the network and estimate the coefficients of their interaction (Kholodenko et al., 2002). Based on the MRA results, we construct a detailed ODE model of the GRN, including all possible regulatory links. We then use the recently developed Bayesian model discovery tool SLING (Jørgensen et al., 2023) to both estimate the parameter values and assess which links are necessary for the gene network to explain and predict experimental data. Taken together, we propose a new network architecture for the core seam cell gene regulatory network.

## Results

### Gene expression analysis via smFISH identifies new interactions between core seam cell TFs

To resolve the interrelationship between three core TFs of the seam cell gene network, namely ELT-1, CEH-16 and EGL-18, we compared gene expression between the wild type and corresponding loss-of-function mutants. Given that both *ceh-16* and *elt-1* are essential genes in *C. elegans*, we chose to utilise hypomorph mutants that were viable, yet displayed a significant decrease in seam cell number, similar to what was observed using a strong *egl-18* loss-of-function allele (Eisenmann and Kim, 2000) (Figure 2A). We used smFISH to quantify mRNA levels in individual seam cells (Figure 2B-C). To ensure cells were behaving similarly, we performed quantification after the L1 asymmetric division, i.e. before the seam cells expanded their number from 10 to 16 via a symmetric division. We chose to focus on V1-V4 lineages as these seam cells are in close proximity along the body axis and behave in a similar manner when it comes to division patterns (Figure 1A). Also, the expression of *elt-1, egl-18* and *ceh-16* follows a similar pattern in V1-V4 seam cells, which is evident from the high correlation in the expression of these genes in V1-V4 cells across different worms in all genetic backgrounds (Figure S1). We found that *elt-1* mRNA levels showed no change in both *ceh-16(bp323)* and *egl-18(ga97)* mutants compared to wild type (Figure 2C). Furthermore, *egl-18* expression was significantly higher in the *ceh-16(bp323)* mutant, while it was similar to the wild type in the *elt-1(ku491)* as well as the *egl-18(ga97)* mutant (Figure 2C). Instead, *ceh-16* expression was decreased in the *ceh-16(bp323)* mutant and even more so in the *egl-18(ga97)* and *elt-1(ku491)* mutant backgrounds (Figure 2C). Taken together, we conclude that CEH-16 is likely a repressor of *egl-18* expression and that *ceh-16* expression may require input from both ELT-1, EGL-18 as well as CEH-16 through a potential self-activation loop.

**Figure 2.**
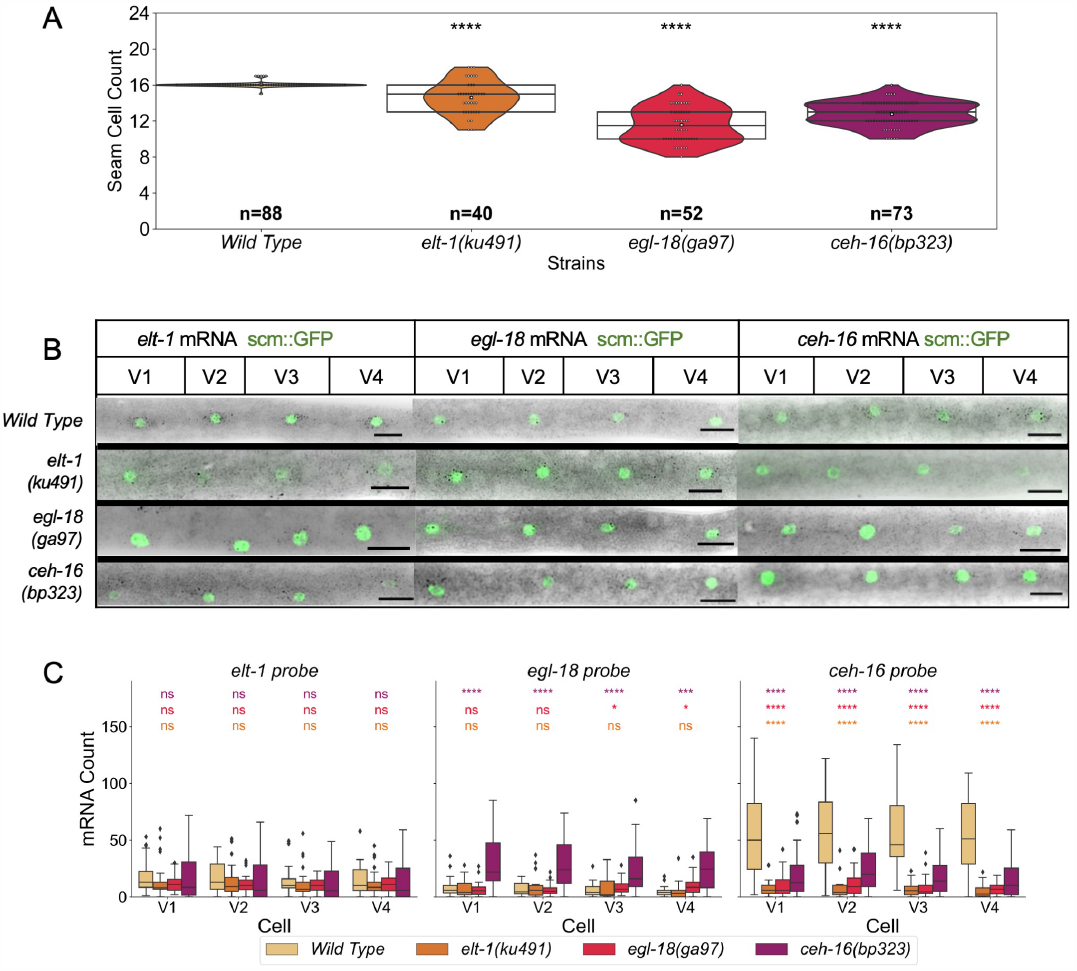
Phenotypic and gene expression analysis in wild type and TF mutant backgrounds. **A:** Seam cell number comparison at the end of the L4 larval stage between the wild type and animals carrying mutations in core seam cell TFs. The lines of the box represent the quartiles of the data, and the white square in the middle represents the median. The p-values are calculated using the Z-test and are shown as * = p *<*0.05, ** = p *<*0.01, *** = p *<*0.001, **** = p *<*0.0001. The n value on the graph corresponds to the number of animals counted. **B:** Representative smFISH images in wild type and TF mutant backgrounds. The mRNAs are visualised as black spots, and the nucleus of the seam cells appears green because all animals carry scm::GFP in the background. **C:** Quantification of smFISH signal in V1-V4 seam cells. In (A) and (C), the significance of the changes between mutant and wild type backgrounds was calculated using the Mann-Whitney U test and is presented as follows: * = p *<*0.05, ** = p *<*0.01, *** = p *<*0.001, **** = p *<*0.0001. Strains are colour-coded as indicated in the key underneath panel C.

We note that gene expressions for these three genes show high levels of worm-to-worm variability characterized by Fano factors well above 1 (Figure S2A), which is the baseline variability for a Poisson process. The large variability was also illustrated by the coefficient of variation (Figure S2B) which was between 0.6 and 1.2 for all the data. While this could be indicative of bursty gene expression, high correlations between the expression of the same gene in the neighbouring cells in the same worm (Figure S1) suggested the presence of large worm-to-worm extrinsic variability. This extrinsic variability can mask the information contained in the intrinsic noise. Therefore, in this study, we mainly focus on the effects of the perturbation on the mean expression.

### Constructing a gene regulatory network using Modular Response Analysis

The smFISH observations were not fully compatible with the naïve network in the literature consisting of a positive action from CEH-16 on *egl-18* and from ELT-1 on *ceh-16* and *egl-18* (Thompson et al., 2016). Therefore we used mathematical modelling and statistical inference to deduce the topology of the underlying GRN by characterising the strength and importance of all new interactions. First, we carried out MRA to estimate the strengths of putative links between the genes of the network (Kholodenko et al., 2002)(see Methods for more details). The MRA algorithm was originally designed for cases where mutants were defined by a change in the quantity of available protein, for example, upon RNAi knockdown. We therefore adapted the MRA algorithm for use in cases where protein activity may be decreased without changes in protein levels, as expected in the case of hypomorph mutants. The MRA produces an interaction matrix where the value of each column indicates the gene carrying out the action, the row indicates the gene receiving the action, and the cell at the overlap between the row and column indicates the strength of the interaction and whether it is activatory (positive) or repressive (negative).

The MRA algorithm was applied to bootstrapped smFISH data collected from V1-V4 cells, and thus a range of coefficients was generated (Figure 3A). The resulting interaction matrix was obtained by taking the average coefficient values, thereby generating a summary GRN (Figure 3B). Strong connections in this case are the positive effect of ELT-1 on *egl-18* and *ceh-16* expression, the negative effect from CEH-16 on *egl-18* expression, and the positive effect from EGL-18 on *ceh-16* expression. A further repression from CEH-16 on *elt-1* and an activation from EGL-18 on *elt-1* are less robust (confidence intervals of 95 % include the value 0, see Figure 3A). These conclusions were also supported when all seam cells were included in the analysis as opposed to V1-V4 only (Figure S3). Taken together, EGL-18 and CEH-16 appear to act downstream of ELT-1, which would be in line with previous observations (Huang et al., 2009; Koh and Rothman, 2001), while CEH-16 and EGL-18 engage in a negative feedback loop that has not previously been described.

**Figure 3.**
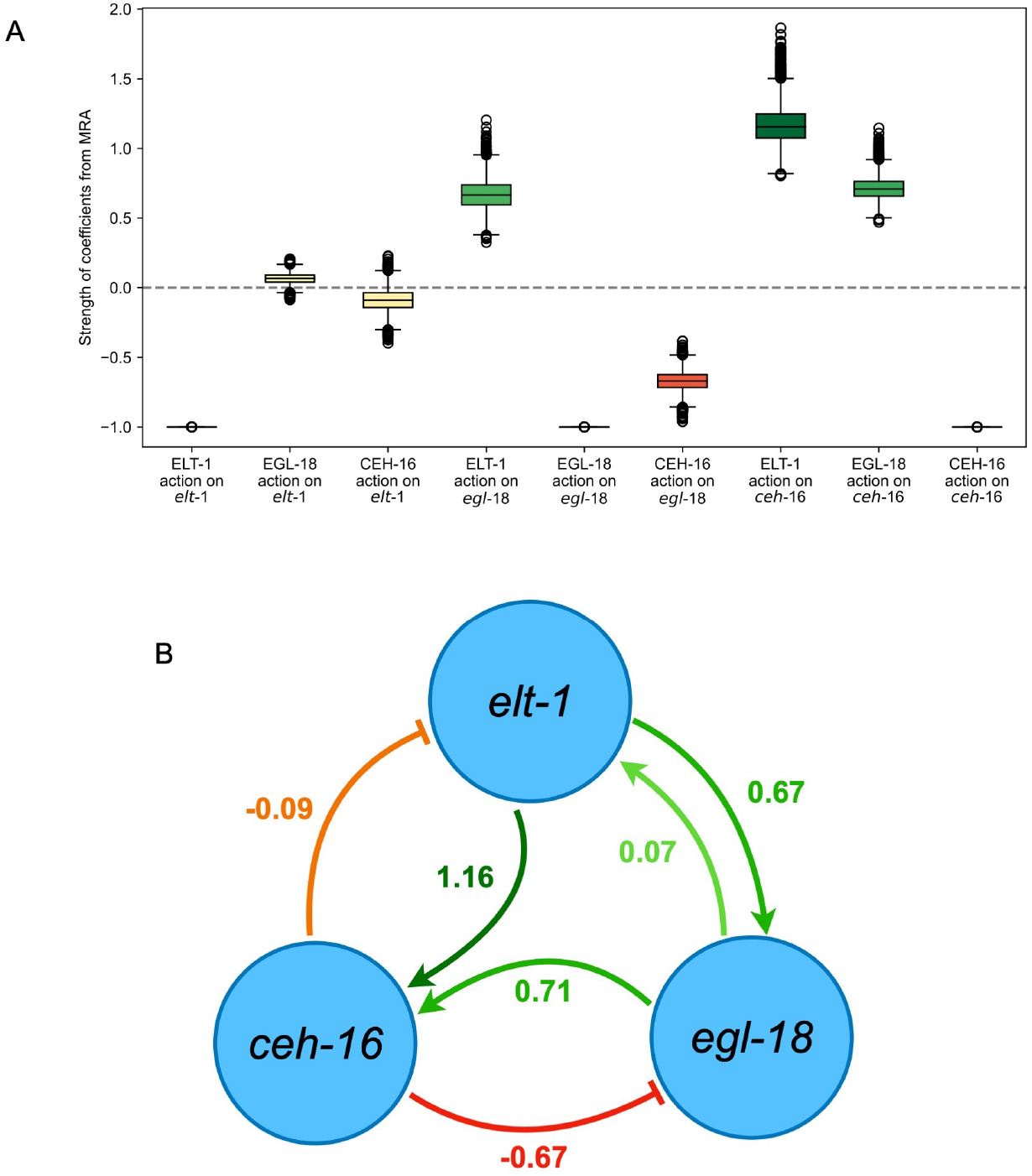
Modular Response Analysis (MRA) to construct a seam cell gene network. **A:** Boxplots of network interaction map coefficients from the MRA performed on the bootstrapped smFISH data of V1-V4 seam cells. A positive number indicates activation, a negative number repression, and 0 indicates a lack of an effect. The coefficients of a gene acting on itself are set to -1 as default in the MRA algorithm. **B:** Network summarising the results of the MRA algorithm using the average of each coefficient value shown in the boxplots in panel A. Green arrows represent activation, while orange/red blunt-end arrows represent repression.

### Inferring an ODE model explaining the gene expression data

To further evaluate the significance of each connection in the regulatory network and derive a model with the least possible number of interactions, we produced an ODE-based description of the wild type and mutant gene expression data (see Methods for details). We defined positive MRA numbers as activation actions and negative numbers as repression actions. We also included a self-activation loop for *ceh-16* as the data indicated a strong loss of *ceh-16* expression in the *ceh-16(bp323)* mutant background. However, the MRA is inherently incapable of detecting self-regulatory loops. When multiple genes regulated another one, we assumed all the possible types of co-regulation in the form of logical OR and AND actions (Table S2A). For example, in the case of only OR actions, the resulting ODE model is:

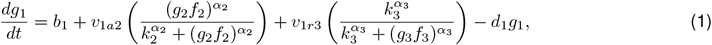

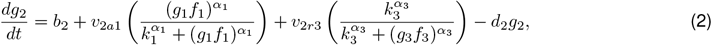

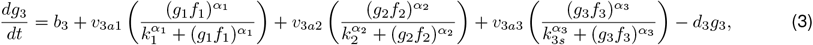

where

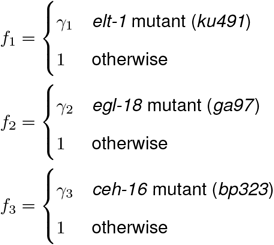

Here, *elt-1* is identified as gene 1 (*g*_1_), *egl-18* as gene 2 (*g*_2_) and *ceh-16* as gene 3 (*g*_3_). The derivatives of their quantities in time are described by equations (1), (2) and (3), respectively. The full list of parameters used in these equations is shown in Table S1A. Degradation rates *d*_1_, *d*_2_ and *d*_3_ were set to 1 since we are not modelling the time evolution of the system, and we make the assumption that the cells at the late L1 stage are at steady-state for the purposes of our ODE model. Activation and repression actions are modelled using Hill functions, as described in the Methods. The functions *f*_1_, *f*_2_, *f*_3_ represent the fraction of protein that is fully functional with 1, 2, 3 referring to ELT-1, EGL-18 and CEH-16, respectively. In ODE modelling of GRNs, co-regulation by logical OR is specified as Hill functions appearing as additive terms in the equation. In contrast, co-regulation by logical AND is specified as Hill functions appearing as multiplicative terms in the equation. We named this system of equations that assumes all actions are linked with the OR relationship (i.e. all are independent and additive) as model 1. From this model, we generated another 19 types of GRN ODE models, where the OR relationships between actions are changed to ANDs in all possible combinations giving rise to 20 model variants described in Table S1B.

These models have all possible regulatory links suggested by MRA and our data, and they include up to 11 unknown parameters. To constrain the parameters of our 20 model variants to data, we use the recently developed Bayesian model discovery approach, SLING. In short, SLING attempts to find the simplest model that explains the data by fitting parameters of the model, setting parameters associated with regulatory links not required to explain the data to zero (see Jørgensen et al., 2023, and Methods for more details). This approach helps to avoid overfitting complex models and encourages simpler models that can explain the data. In our case, SLING is particularly useful to test which gene interactions are necessary to describe the system in the wild type and mutant settings. Note that this approach highlights whether these interactions are necessary in order to fit the data within a reasonable margin of error, yet it does not rule out the actual presence of some non-essential links. SLING is a simulation-based inference method and relies on a distance measure between the summary statistics of the data and the model outputs. In this case, we are using the mean expression of our 3 TFs of interest in V1-V4 seam cells in the wild type and mutants (shown in Figure 2C) as the summary statistics of the data. Model predictions are the numerically obtained steady-state solutions of the ODE for each choice of parameters, and we use a normalised Euclidean distance between the data and model outputs (see Methods for details).

We allowed the model to fit the parameters of type *b, v* and *k*. We set the degradation parameters to 1 and *γ*_2_ = 0.01 and *γ*_1_ = *γ*_3_ = 0.25 to match the predicted nature of the genetic alleles with almost complete loss for EGL-18 and partial loss for ELT-1 and CEH-16 function. We performed 3 independent Markov Chain Monte Carlo runs using SLInG for each of our 20 model variants. We then calculated how frequently the parameters were present in the final model fits (Figure 4A), from which we also derived the importance of links and their relationships (Figure 4B). For example, the basal production of *elt-1*, that is, the production of mRNA that is independent of the levels of the other genes in the network, is highly present (97%) in the model fits. However, the basal production of *egl-18* and *ceh-16* is only present in 18% and 0% of the model fits, respectively. This indicates that *ceh-16* expression and, to a lesser extent, *egl-18* expression rely highly on the presence of the two other core TFs while *elt-1* expression appears to be largely independent.

**Figure 4.**
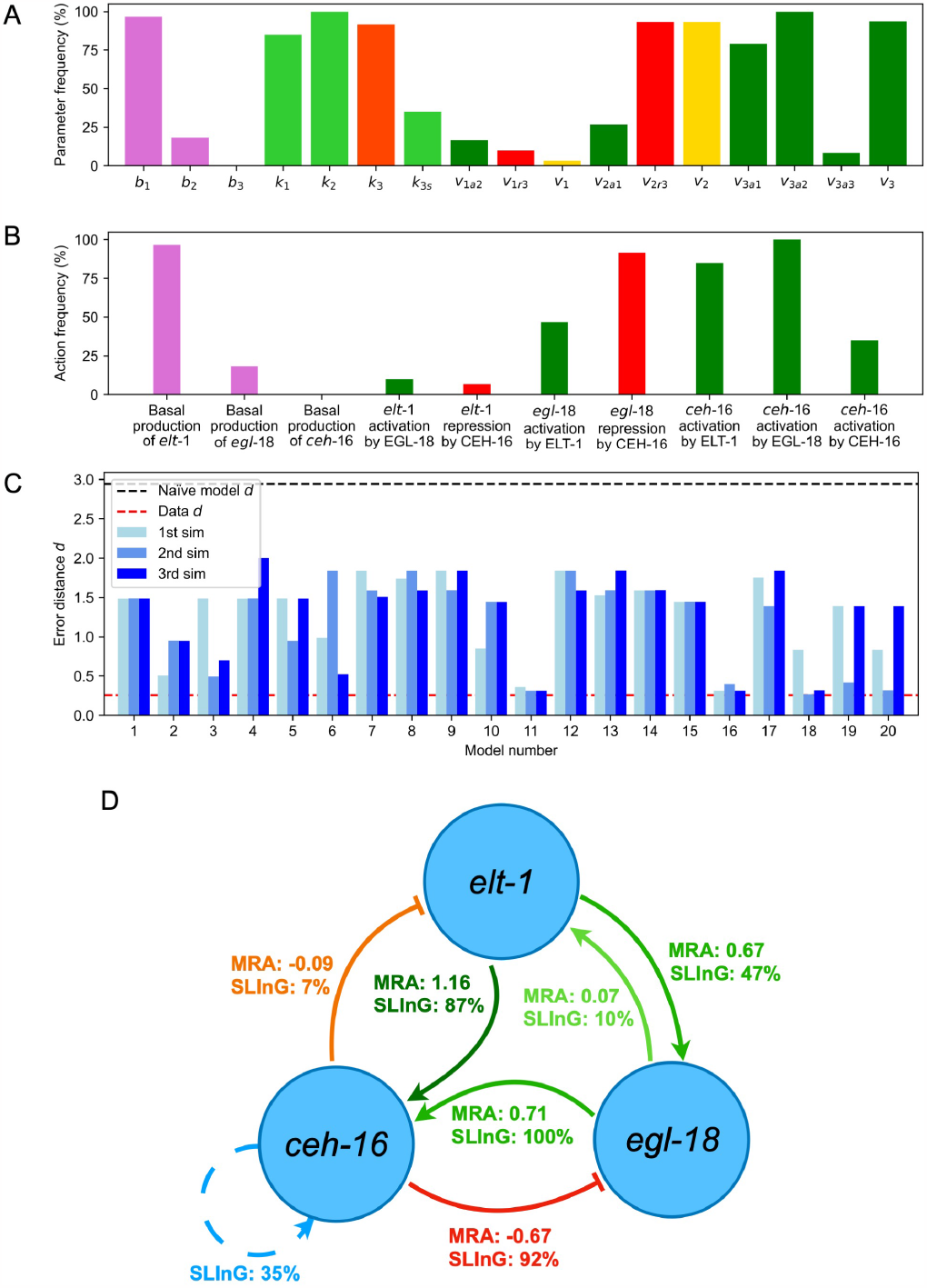
Model fitting using SLING. **A:** Bar graph showing the presence of each parameter in the final models, computed as the percentage of all SLInG runs. **B:** The presence of each effect (in per cent) in the final models is shown. These are derived from the combinations of parameter presence in each SLInG run. **C:** Distance measure of the median parameter set d, grouped by model type (3 SLInG runs for each). The red line denoting the data value indicates one standard deviation of the original data. **D:** GRN summarising the MRA and SLInG results. The percentages for the SLInG reflect the presence of actions in final models.

We compare 3 SLInG runs of the 20 models (blue bars) to the data (red dashed line) using the ABC distance measure(Figure 4C). We also compare our model and the effective data to the naïve model, also using the ABC distance measure (black dashed line) (Figure 4C), which refers to the model with only activation actions taken from the literature (Figure 1B). All our models performed better than the naïve model, with some SLInG runs being very close to the data distance (this is the closest we expect the model to be with the data given the standard deviation of the bootstrapped data). If we consider the final model fits of all model variants as equal, we can calculate the frequency in which each interaction appears (Figure 4D). Both the CEH-16 and EGL-18 effect on *elt-1* is rarely present (10% and 7% respectively). The *egl-18* activation by ELT-1 is kept in around half the cases (47%). However, this link is mostly present when linked with an AND to the *egl-18* repression by CEH-16, which is present in 92% of the cases. The *ceh-16* activation by ELT-1 is also highly present (87%) and *ceh-16* activation by EGL-18 is always maintained (100%). The *ceh-16* self-activatory loop is there in some SLInG runs (35%) (Figure 4C).

Next, to assess whether there is high cooperativity in the binding of the core TFs, we compared the model parameters trained with *α* = 1 (no cooperativity between proteins) as opposed to *α* = 3 (cooperativity). Here, we operate 3 SLInG runs for the two extreme models, either with all actions being OR (model 1) or AND (model 20), resulting in a total of 6 SLInG runs. These SLInG runs revealed no significant differences when focusing on the frequency of parameters representing the network interactions (Figure 5A). The distances of the SLInG runs of the two models (Figure 5B), were also very similar. While we could have also fitted the *α* exponents, we decided to fix *α* = 1 in our main SLInG runs, as the chosen gene network does not significantly depend on the exponents of the Hill function, and thus whether there is cooperativity or not.

**Figure 5.**
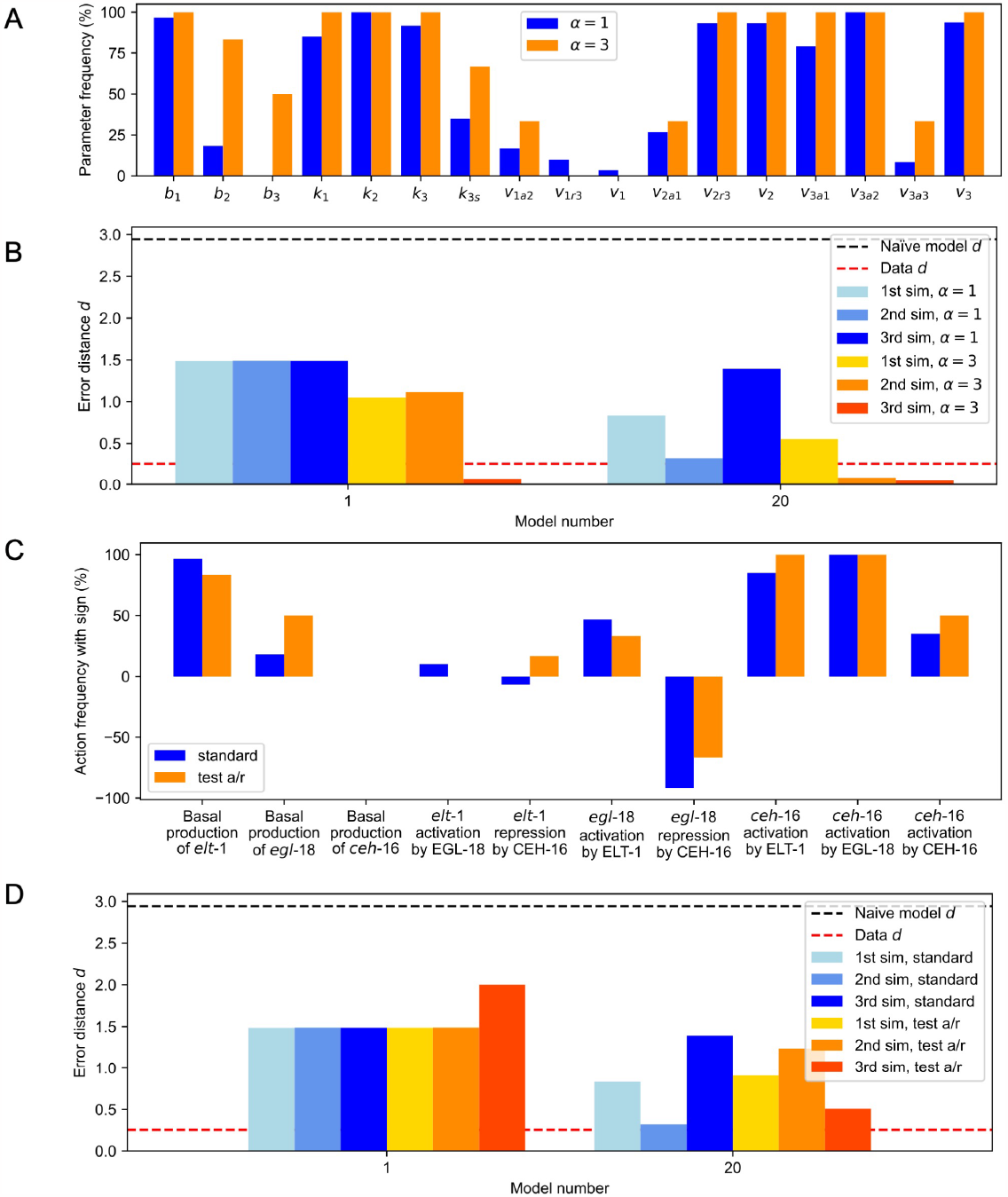
Testing additional conditions for SLING: **A:** Bar graph showing parameter presence in the final models comparing α = 1 (60 SLInG runs, 3 for each one of the 20 models) with α = 3 (6 SLInG runs, 3 for models 1 and 20). **B:** Distance measure of the median parameter set in the SLInG runs for models 1 and 20 comparing α = 1 and α = 3. **C:** Bar graph showing the percentage and sign of the actions of the standard model compared to the activation/repression (a/r) model. **D:** Distance measure of the median parameter set of the standard model compared to the activation/repression (a/r) model.

We next decided to relax the assumption on the type of regulation we have observed from the MRA and let SLInG test both activation and repression (denoted here by a/r) for every single link and decide which one is best to keep or whether both can be discarded. To achieve this, we gave all *v* parameters the option to be positive, negative or 0. If a parameter is kept, then the action depends on its sign, and its absolute value is used as the rate. We selected models 1 and 20, again, to carry out this analysis. The comparison of these new models with the standard SLInG runs revealed a very similar behaviour to the standard models(Figure 5C and 5D). Taken together, these results reinforce the results obtained by the MRA and SLInG.

### Model validation using double mutant expression data

The ODE GRN model allows us to generate predictions as to what the behaviour of the system would be upon the removal of two nodes at once. Given that an unexpected outcome from the model was the negative feedback loop that involved CEH-16 mediated repression of *egl-18* and the EGL-18 driven activation of *ceh-16*, we decided to construct a *ceh-16(bp323);egl-18(ga97)* double mutant and test model predictions against new experimental data. First, we compared the double mutant with the two parental strains phenotypically by carrying out seam cell counts at the end of the L4 larval stage. We found a significant decrease in seam cell number in comparison to the wild type in both parental strains and the double mutant. We then compared the double mutant to the parental strains and noted both a decrease in mean seam cell number and variability of seam cell counts(Figure 6A). *elt-1* mRNA levels did not change significantly from the wild type in the *ceh-16(bp323);egl-18(ga97)* mutant. Instead, we found that *egl-18* levels show a significant increase compared to the wild type and that *ceh-16* levels decrease (Figure 6B).

**Figure 6.**
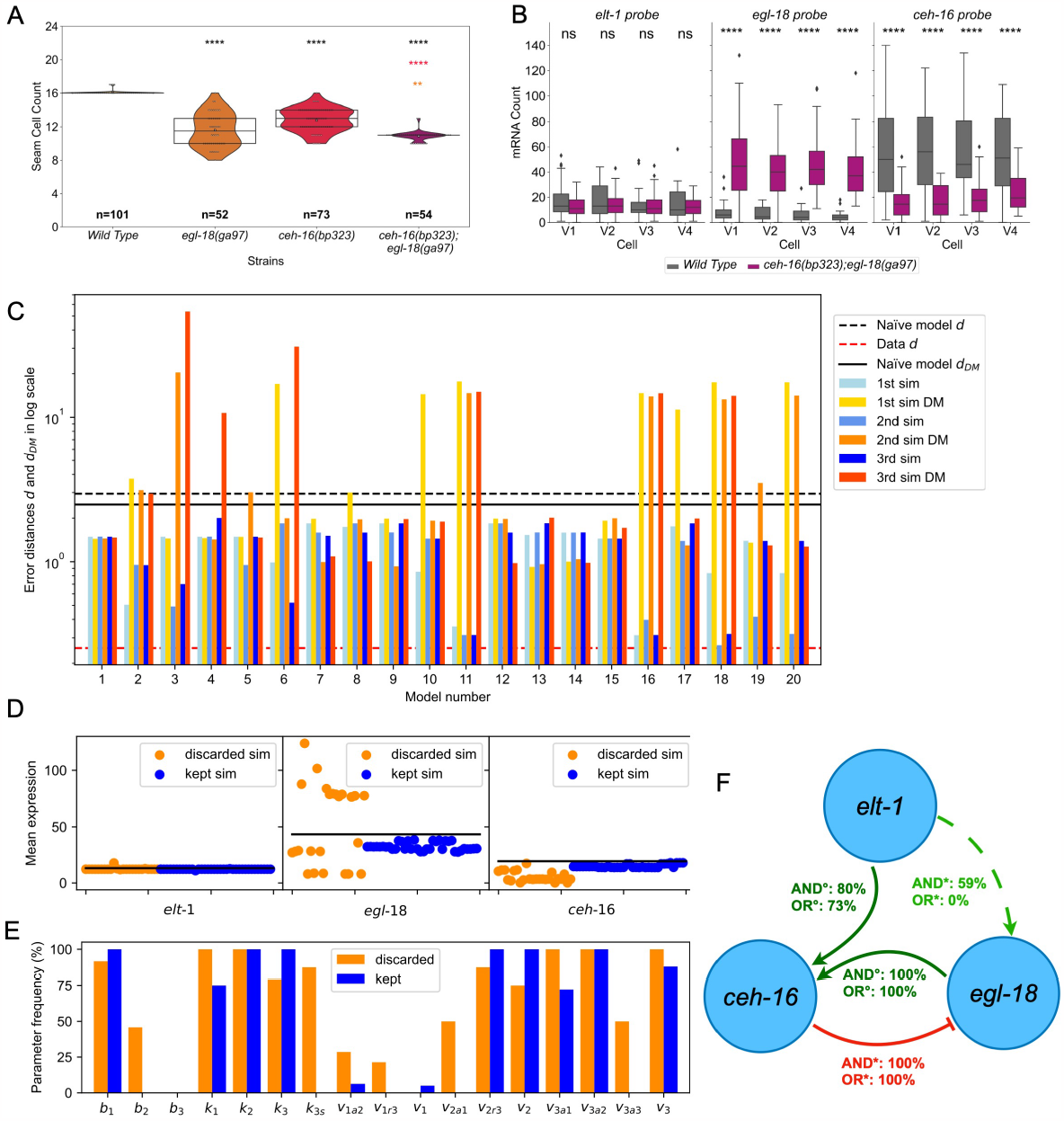
Figure 6: Validating model predictions using a double mutant. The significance of the changes in cell number was calculated using the Z-test and denoted as * = p *<*0.05, ** = p *<*0.01, *** = p *<*0.001, **** = p *<*0.0001. The comparison between all the mutants and wild type is denoted in black asterisk, and the comparison between the double mutant and the parental strains is denoted with asterisks in the colour of the corresponding parental strain. We additionally calculated the difference in variability using the Levene’s test between the double mutant and wild type, *ceh-16(bp323)* and *egl-18(ga97)* and the resulting p-values were 1.85 x 10^−4^, 3.35 x 10^−10^, and 2.37 x 10^−16^ respectively. **A:** Seam cell counts in the double *egl-18(ga97); ceh-16(bp323)* mutant in comparison to the wild type and the parental strains. **B:** Quantification of *elt-1, egl-18* and *ceh-16* expression in V1-V4 seam cells via smFISH in the wild type and *egl-18(ga97); ceh-16(bp323)* double mutant. The significance of the changes in expression was calculated with respect to the wild type using the Mann-Whitney U test and denoted as * = p *<*0.05, ** = p *<*0.01, *** = p *<*0.001, **** = p *<*0.0001.**C:** Distance measure *d* (on a logarithmic scale) of the median parameter sets in predicting the wild type and single-mutant behaviour vs the double mutant behaviour. The bars are superimposed with the data presented in Figure 4C, with the lowest distance measure (depending on the SLInG run) shown in front. **D:** Predicted gene expression in the double mutant from SLInG runs (using the median parameter set) is compared to data (black lines). SLInG runs are grouped into 2 categories based on their double-mutant error: they are discarded when 2.5 *<*d and kept when *d* ≤ 2.5. **E:** Parameter presence (in % of models) for the two categories of SLInG runs is shown. **F:** Final model based on all kept SLInG runs that are able to predict the double mutant data. The possible interacting actions on *egl-18* are indicated with the symbol *, while the activations on *ceh-16* are indicated with the symbol °. The percentage shown on each link is the number of times each connection is kept given an initial model that supposes an AND/OR relationship with the effect on the same gene by the other factor.

We next tested our model predictions using these results. All ODE models produced good fits when tested on single-gene mutations, so these were subsequently tested with the double mutant data. The model prediction errors for the double mutant data were often higher, with some models performing worse than the naïve model (Figure 6C). Models that performed poorly in the double mutant context (mean expressions with a distance higher than 2.5 in any the simulations) were discarded (Figure 6C-D). Levels of *elt-1* mRNA were accurately predicted in all models (Figure 6D). The mean expression of *egl-18* had the most variable predictions, which was particularly evident in the models we discarded. Furthermore, in all discarded SLInG runs, *ceh-16* expression was underestimated, meaning the best models are the ones with the highest estimate of *ceh-16* levels (Figure 6D).

We also investigated how the parameter selection varies between the two groups of models (Figure 6E). The activation of *ceh-16* by EGL-18 (*v*_3*a*2_) is kept in all SLInG runs independently of whether they are discarded or kept, thus indicating that it’s one of the core principles that allow our model to function. However, some parameters such as the basal production of *egl-18* (*b*_2_) or the *ceh-16* self-activation (*k*_3*s*_) are only kept in models that were discarded in the context of the double mutant data, indicating that including them may result in overfitting. This new analysis allowed us to narrow down the OR/AND logic of some network connections too. For example, the activation of *egl-18* by ELT-1 when linked with an OR logic to the repression by CEH-16 (*v*_2*a*1_) is only kept in the worst fitting SLInG runs. Nevertheless, when linked by an AND, the link was kept in 59% of the cases indicating that this network link can only exist if *egl-18* is also repressed by CEH-16. By deleting the links that are discarded in 90% or more of the cases, we obtained the final network model that reflects the results from the kept SLInG runs (Figure 6F). Based on this, the first equation (for the derivative of *elt-1*) can be described without any regulation by simply using constant synthesis and constant degradation rate:

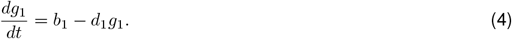

The second equation (for the derivative of *egl-18*) can have two possible forms

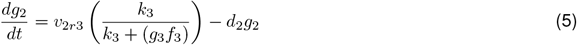

and

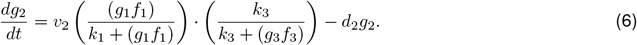

The third equation (for the derivative of *ceh-16*) also has two possible forms

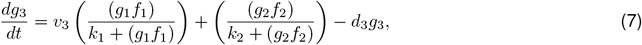

or

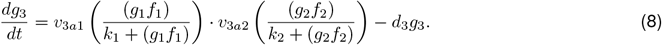

Hence, the ODE models of the final network that best describe our single-mutant and double-mutant data are obtained by combining equation (4), with (5) or (6), and (7) or (8). This network is illustrated in Figure 6F, and it includes an incoherent feedforward network and negative feedback.

## Discussion

Inference of gene regulatory networks is fundamental to increasing our mechanistic understanding of various biological processes, so there is a continuous need to develop approaches to tackle this problem at different levels. Our aim here was to use an integrative experimental and computational approach to revisit a three-factor GRN underlying seam cell development in *C. elegans*. Using smFISH analysis, we were able to quantify mRNA levels in individual seam cells in wild type and mutant backgrounds. While smFISH data and *C. elegans* gene transcription itself can have high levels of noise, we were able to identify clear patterns that were then used to infer the network architecture and extrapolate new interactions from this data using mathematical approaches. The adaptation of the MRA algorithm (Kholodenko et al., 2002) to genetic hypomorphs has given us the flexibility to work without gene knockouts, which was necessary in this case as two out of the three core factors represent essential genes. The MRA produced an initial quantitative description of the network, which we built on by searching among a series of ODE models, using the recently proposed Bayesian model discovery approach, SLInG (Jørgensen et al., 2023). This also allowed us to narrow down which interactions are essential to explain the data resulting in several candidate ODE models using different regulatory logics. Finally, we quantified gene expression from a double mutant to validate our ODE models. This confirmed our models’ ability to predict behaviours of the core TF GRN for seam cells and enabled us to obtain the most likely ODE model describing our system. Combining these approaches, we present an updated core seam cell gene network architecture. This updated network will be valuable to increase the mechanistic understanding of the spatial patterning system and to interrogate the relationship of other components to the core factors.

Gene expression is highly stochastic due to the inherent random timing of biochemical reactions and also extrinsic variability in the cellular environment (Shahrezaei and Swain, 2008). The former type of stochasticity, which is sometimes referred to as intrinsic noise, is highly informative of the underlying regulatory pathway. Indeed, several studies have argued that this type of noise can be harnessed for learning the underlying GRN (Munsky et al., 2009; Stumpf, 2021). However, extrinsic noise caused by global factors in the cell or tissue creates correlations between transcripts and proteins that are independent of the underlying regulatory relationship, masking correlations induced by regulation in snap-shot data (Dunlop et al., 2008). We have observed high levels of expression noise in our data, which is dominated in this case by extrinsic noise. Therefore, here we have decided only to focus on the mean transcription expression and a deterministic (ODE) description of our GRN. In future, one could use methods such as dual-colour smFISH (Kramer et al., 2021), two-colour TF quantification experiments (Elowitz et al., 2002) or time-lapse imaging (Dunlop et al., 2008) to decouple extrinsic and intrinsic noise in the seam cells and their link to the underlying regulatory architecture.

The seam cell gene network that could be derived from the literature included an activation from ELT-1 to *ceh-16* and *egl-18*, and it portrayed CEH-16 as an activator of transcription of *egl-18* (Huang et al., 2009; Cassata et al., 2005; Thompson et al., 2016). Our smFISH and modelling data suggested that *elt-1* expression does not depend on EGL-18 and CEH-16, which is consistent with the key role for ELT-1 as a master regulator in the specification of the epidermis (Spieth et al., 1991; Page et al., 1997). However, we found two new interactions that are consistently present in our models and involve the activation of *ceh-16* by EGL-18 and the repression of *egl-18* by CEH-16. Previous work had shown *egl-18* to be regulated by CEH-16 during embryonic development; however, this interaction had been described as positive based on the observed decrease in expression of an EGL-18::GFP reporter upon *ceh-16* RNAi (Cassata et al., 2005). Instead, we find that CEH-16 negatively regulates *egl-18* in early larval seam cell development, mirroring the more established repressive function of its homolog *engrailed* (Markel et al., 2002; Alexandre and Vincent, 2003; Genestine et al., 2015; Huang et al., 2009). This discrepancy may reflect the different nature of the genetic perturbations used in these studies, namely a *ceh-16* hypomorph in our case, which leads to minimal loss of seam cell fate as opposed to strong *ceh-16* RNAi in the earlier study. Strong loss of CEH-16 function leads to seam cell fusion and high embryonic lethality, in which case loss of seam cell marker expression may not be a direct effect of CEH-16 action on *egl-18* transcription (Cassata et al., 2005). Nevertheless, given that *engrailed* is known to have both repressive and activating functions (Alexandre and Vincent, 2003), we cannot rule out that the fine-tuning of the seam cell network varies depending on the developmental stage. Future studies should include more temporal data to evaluate whether the behaviour of the network remains similar beyond the late L1 stage used in our study. Furthermore, it remains unclear whether the described interactions in our model are direct or not, with modENCODE ChIP-seq data from later larval stages indicating potential direct binding of CEH-16 and ELT-1 on the *egl-18, elt-1* and *ceh-16* locus (Luo et al., 2020; Consortium, 2012; Kudron et al., 2018).

Our genetic analysis revealed an unexpected interaction between *ceh-16(bp323)* and *egl-18(ga97)* mutants, with the double mutants showing a significant decrease in phenotypic variance in relation to the parental strains. This result is surprising because seam cell mutants tend to be generally variable, although in this case, the double mutants restore developmental robustness to comparable levels to wild type animals. We note that this interaction is compatible with the updated network architecture as it is consistent with the feedback loop proposed between EGL-18 and CEH-16. We hypothesise that double mutants may show stronger failure of symmetric divisions at L2 in comparison to *ceh-16(bp323)* animals because the EGL-18 activation input is lacking. In addition to this, seam cell fate acquisition may be strengthened in the double mutants due to the lack of repression of *egl-18* by CEH-16. Given that *egl-18(ga97)* is a nonsense mutation, the strengthening effect on seam cell fate is likely to be driven in the double mutant not directly by EGL-18 but through other factors that promote seam cell fate, with one candidate being its close paralogue ELT-6 (Koneru, 2020; Koh et al., 2002; Koh and Rothman, 2001).

We note that our inferred gene regulatory network contains negative and possibly positive feedback and also an incoherent feed-forward loop. Depending on network parameters, such systems could exhibit a range of dynamic properties that could underlie the robust cell fate decision-making. Studies of cellular decision-making from different organisms suggest a core regulatory network structure consisting of multiple nested feedback loops, exhibiting specific nonlinear dynamical behaviours such as bi-stability or excitability required for robust cell fate determination (Balázsi et al., 2011; Moris et al., 2016; Sáez et al., 2022; Petratou et al., 2018). Our study adds to this body of work by revealing the architecture of the core gene-regulatory network in epidermal stem cells of *C. elegans*. Future experimental and computational characterisation of the dynamical properties of this system could elucidate the link to the observed developmental robustness.

## Methods

### C. elegans strains and culture

Nematodes were cultured according to standard protocols (Brenner, 1974). All nematodes were grown on NGM plates with OP50 as a food source. As reference strain, we used the JR667 strain, which contains an *scm::GFP* transgene used to visualise the seam cells *(wls51)*. Mutant strains used in this study were MBA650(*elt-1(ku491) IV; wIs51[scm::GFP + unc-119(+)] V)*, MBA290*(egl-18(ga97) IV; wIs51[scm::GFP + unc-119(+)] V)*, HZ620*(ceh-16(bp323) III; wIs51[scm::GFP + unc-119(+)] V)* and MBA1216*(ceh-16(bp323)III; egl-18(ga97) IV* ; *wIs51[scm::GFP + unc-119(+)] V)*.

### Single Molecule Fluorescent In Situ Hybridisation (smFISH)

The synchronised population subjected to smFISH was fixed at 17 hours post placing eggs on an NGM plate. This timing corresponds to the late L1 stage, as also confirmed via microscopy. Imaging and smFISH analysis was performed as previously described (Hintze et al., 2020). Images were taken using a motorized epifluoresence Ti-eclipse microscope (Nikon) and a DU-934 CCD-17291 camera (Andor Technology) in 0.8 μm interval z-stacks using a 100x objective. mRNA spot quantification was carried out using MATLAB (MathWorks) as previously described (Barkoulas et al., 2013). The images were processed through ImageJ (NIH, Rockville, MD). The probes used are shown in Table S2.

### Mathematical Modelling

#### Building an ordinary differential equation (ODE) model from a GRN

GRNs are suitable to represent the interaction of TFs, which are proteins (denoted below in capitals, such as GENE 1), with the promotors to control transcription of mRNAs (denoted below in italics, such as *gene 1*). As transcription is typically much faster than translation, we can use time-scale separation and only write ODEs for one set of variables, assuming mRNA levels and protein levels are proportional to one another. As we have data for mRNA levels, we are assuming our variables represent mRNA levels. We describe below the detailed steps to build an ODE model to describe a GRN from its directed graph representation:

1. Create an ordinary differential equation for each *gene i* present in the network, starting by

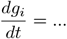

where *g*_*i*_ describes the level of mRNA of *gene i* over time (*t*).
2. Start building the right-hand side (RHS), adding a basal term of production *b*_*i*_ and subtracting a degradation term *d*_*i*_*g*_*i*_, where *d*_*i*_ is a degradation rate. So, now,

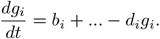
3. Consider all the GENES that regulate *gene i*. For example, consider the case where GENE J activates *gene i* and GENE L represses it; in this scenario, we incorporate two functions to the RHS (*H*_*ial*_(*g*_*j*_) and *Ĥ*_*irl*_(*g*_*l*_)) that represent these actions respectively (e.g. *ial* is read as ‘*gene i* is activated by GENE L’). Hence, the equation becomes

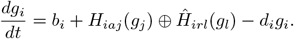
4. Substitute the ⊕ with either a sum (+) or multiplication (*·*), depending on whether the actions of J and L in the network are related with an OR (+) or an AND (*·*).
5. The activation of *gene i* by the action of GENE J is described by the Hill–Langmuir function

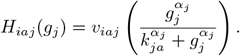

Here, *v*_*iaj*_ is the maximum activation rate of *i* by L, which is multiplied by the Hill function 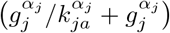. The coefficient *k*_*ja*_ is the quantity of *j* to produce half occupation, i.e. the quantity of *j* that gives the fraction a value of 1/2 (Nelson et al., 2008). The exponents *α*_*j*_s are positive numbers: When greater than 1, they represent a certain degree of cooperativity amongst genes J binding as activators, while when they are 1, they indicate no cooperativity (Klipp et al., 2005).
6. The repression of *gene i* by GENE L is described by the Hill–Langmuir function

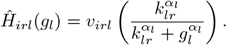

Here, *v*_*irl*_ is the maximum repression rate of *i* by L, which is multiplied by the Hill function 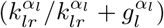. The coefficient *k*_*lr*_ is the quantity of *l* to produce half occupation, i.e. the quantity of *l* that gives the fraction a value of ^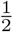^ (Nelson et al., 2008). The exponents *α*_*l*_ represent the degree of cooperativity amongst genes of type L binding as repressors (Klipp et al., 2005).
7. In case a *gene i* self-activates (*i* is activated by its protein I), a function

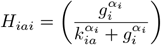

should be included. In the case of self-repression, Ĥ_*iri*_ should be included. The self-action term should be added to the other actions if linked with an OR, or multiplied if linked with an AND.
8. Note that all parameter constants are non-negative. A zero *v* indicates the lack of that action. A zero *k* in a repression action sets the whole fraction to 0 when the denominator is non-zero, while in an activation action, it sets the fraction to 1. When two effects are multiplied (AND), their *v*s can be substituted by one unique *v*.
9. The coefficients *k* allow the direct use of the mRNA mean expression in the equations, rather than the protein level, under the assumption that these two are proportional. Indeed, if we wanted to use the protein level *G*_*i*_ instead of *g*_*i*_ in the actions, where *G*_*i*_ = *μg*_*i*_, then we could substitute *k*_*i*_ with *K*_*i*_ = *μk*_*i*_.
10. From this system of ODEs used to describe the wild type data, we can obtain models to describe the mutants’ data by modifying all the Hill-Langmuir functions. The activation proposed becomes

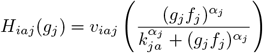

since the quantity of protein J is now *G*_*j*_ = *μf*_*j*_*g*_*j*_. The repression becomes

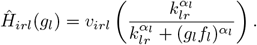

The function *f*_*j*_ (and, similarly, *f*_*l*_) represents a fraction in [0, 1]. The fully-working proteins J produced in the selected genotype over the quantity produced in the wild type, with the same amount of gene *j* mRNA. This function attains a value of 1 in the wild type since all proteins are working. Hence, their binding action is preserved, while they attain the genotype-specific value *γ*_*i*_ ∈ [0, 1) when a mutation of gene j is considered. Note that the value of *γ*_*i*_ can depend on the mutation’s genotype.

As an example, let’s now consider the following network of 3 genes: Gene 1 is not influenced by the presence of Gene 2 and Gene 3. Gene 2 is activated by Gene 1, Gene 2 is repressed by Gene 3, and these two actions are linked by an OR. Gene 3 is activated by Gene 1 AND activated by Gene 2. Gene 3 also self-activates, with an OR link with the other two activation actions of Gene 3. The production of mRNA of *gene 1* is not influenced by the expression of the proteins of GENE 2 and GENE 3; hence, we can write

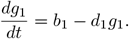

We now consider the expression of *gene 2*, which is activated by GENE 1 and repressed by GENE 2. In the network, we are not specifying any link between those two actions; hence, we consider it an OR link. Thus the equation is

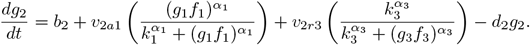

Regarding the gene expression of *gene 3*, we have activation by GENE 1 AND GENE 2, plus a self-activation, linked to them with an OR. Thus the equation is

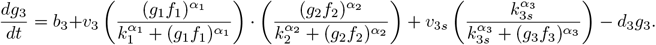

Since GENE 1 is only an activator, we identified *k*_1*a*2_ = *k*_1*a*3_ = *k*_1_. Moreover, we called *k*_2*r*3_ = *k*_2_ but kept two different *k*s for *k*_3*r*2_ = *k*_3_ and *k*_3*a*3_ = *k*_3*s*_ since GENE 3 is a repressor and a self activator.

This system of 3 equations is an ODE model for the GRN. Some values can be assigned to the constants based on experimental estimates or on prior knowledge. Moreover, parameter inference algorithms can be used to find suitable parameter sets that fit the experimental data. We note that while the GRN ODE models can describe transient and temporal dynamics of gene expression in time, in this study, we are assuming that gene expression has reached a steady state at the single developmental time-point the experimental data has been obtained. So, we numerically solve for the steady-state value of expression of different genes while fitting our ODE models to the data.

### ABC distance measure

In order to fit the ODE models, we define an error distance measure *d* between the data for *n* genes and *m* phenotypes used and model output. This is the common practice in likelihood-free inference methods such as Approximate Bayesian Computation (ABC) (Beaumont, 2019), hence we refer to this as ABC distance measure or error distance measure. We denote the mean expression of the data with *μ*_*i,j*_, where *i* ∈ *{*1, 2, …, *n}* indicates the gene and *j* ∈ *{*1, 2, …, *m}* indicates the phenotype (wild type or mutant), and their standard deviation with *σ*_*i,j*_; both are computed using bootstrapping. We also denote the estimated fixed points as (*s*_1,*j*_, *s*_2,*j*_, …, *s*_*n,j*_), for each phenotype *j* as the limit of the ODE simulation and the matrix containing the approximate fixed points information with *S* = *{s*_*i,j*_*}*_*i,j*_. In this notation, the distance measure is defined as follows:

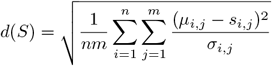

For the main inference task, we had *n* = 3 and *m* = 4, but in the double-mutant testing case, we only used the double-mutant data; hence, *m* = 1.

### Algorithms

#### Modular Response Analysis

Following (Kholodenko et al., 2002), we implemented a Modular Response Analysis (MRA) algorithm, adding a factor that measures the change in the fully working protein production due to gene mutations. For partial-loss-of-function mutants, we have estimated the activity level at 25% due to the fact that the heterozygote behaves like a wild type animal, and thus the function of the gene should be below 50 % but above 0%. For the complete loss-of-function mutants, we estimated the function at 0.01%.

The MRA quantifies the intensity of the influence of a module on another module. These quantifications are measured as perturbations from the steady state (wild type genetic equilibrium) to each mutant state. A module is represented by the synthesis and degradation of mRNA and the mRNA levels. The original algorithm measures the influence of the mRNA level of each gene on the synthesis of the other genes. We consider a module to be a gene’s mRNA level, controlled by its synthesis and degradation, and its protein levels, which we consider to be equivalent to the mRNA mean expressions, except in mutants. Note that, for the purpose of the algorithm, assuming the same amount of protein and mRNA of a certain gene in the wild type is equivalent to assuming a proportional amount with a constant rate since the constant is simplified when considering the ratios.

The MRA algorithm produces a network interaction map (or matrix) *r*, whose elements estimate the type and strength of each action. Each element of the matrix *r*_*ij*_ is called the local response coefficient, and it describes the effect on gene *i* (row) given by mutant *j* (column) (Kholodenko et al., 2002). A positive coefficient indicates an activation action, while a negative one indicates repression. The modulus of each coefficient, |*r*_*ij*_|, represents the strength of the action:

- If |*r*_*ij*_| = 1, the relative changes in *j* are reproduced as equal relative changes in *i*.
- If |*r*_*ij*_| *>* 1, the changes in *j* are amplified in *i*.
- If 0 *<* |*r*_*ij*_| *<* 1, the changes in *j* are attenuated in *i*.
- If |*r*_*ij*_| = 0, *j* has no significant action on *i*.

We apply the MRA algorithm repeatedly using bootstrapping. A single iteration computes the interaction matrix, and all coefficients obtained, shown in Figure 3A. These are then averaged to compute the final interaction matrix, shown in Figure 3B. The mRNA mean expressions (for all genes and all genotypes) are recomputed for every bootstrap sample, picking the data using random sampling with replacement.

The steps we follow to obtain the network interaction matrix (*r*) for each bootstrap sample are as follows (for the full mathematical derivation, see (Kholodenko et al., 2002)). Firstly, a response matrix is computed:

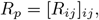

Where

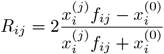

and

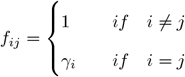

Here, 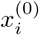 is the mean expression of gene *i* in the wild type, while 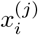 is its mean expression in the mutant *j. R*_*ij*_s are called central fractional differences and are computed as the finite difference of the transcript relative levels 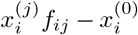 divided by their mean value 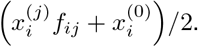. Subsequently, the network interaction matrix is computed:

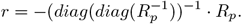

This formula for *r* is based on the assumption that all *r*_*ii*_ = −1; thus, the presence of gene *i* above some level prevents a further increase of this gene expression. This corresponds to the gene’s own degradation, which depends on its level. However, MRA is not able to infer other types of self-loops.

#### SLInG

We have constructed the most general ODE models based on the MRA results. We then employed SLInG (Sparse Likelihood-free inference using Gibbs, Jørgensen et al., 2023) to obtain the simplest possible submodels that can explain the data: SLInG is a Bayesian model-discovery tool based on a Gibbs sampling algorithm and can be used for model selection and parameter inference among a large set of related models. SLInG can be thought of as a sparse Approximate Bayesian Computation (ABC) method (Beaumont, 2019) and builds on the ABC algorithm presented by Turner and Van Zandt (2014) by using sparsity-inducing priors. In Jørgensen et al. (2023), self-consistent tests based on synthetic data demonstrate that SLInG can reliably reverse-engineer GRNs from data and SLInG outperform state-of-the-art methods that rely on constructing GRNs from expression correlation networks.

SLInG performs sparse sampling, which encourages the algorithm to discard parameters that are not needed to explain the data. Different schemes of this type have successfully been applied for equation discovery in recent years (Brunton et al., 2016; Alahmadi et al., 2020; Both et al., 2021). Within a Bayesian setting, sparsity is accomplished by including, e.g., Laplace priors for each parameter, and we take this approach in our analysis with SLInG (Park and Casella, 2008; Chen et al., 2011; Mallick and Yi, 2014, cf. Bayesian LASSO). The sampling scheme of SLInG is hierarchical in the sense that each parameter is associated with a hyperparameter that sets the respective level of sparsity (Davies et al., 2017). SLInG autonomously calibrates these hyperparameters as it explores the parameter space. The hyperparameters are thus iteratively adjusted such that the use of superfluous parameters is highly discouraged while the algorithm is able to properly explore the parameter space along the dimensions that are necessary to describe the data. SLInG hence has the advantage that it can tailor the level of sparsity for each individual parameter of the model without introducing a corresponding number of free hyperparameters to be set by the user.

SLInG introduces an additional sparsity parameter (*δ*_ABC_) that sets the overall level of sparsity by scaling the distance measure, i.e. by changing the weight that the acceptance probability attributes to changes in the likelihood (cf. Turner and Van Zandt, 2014). For the analysis presented in this paper, we set *δ*_ABC_ such as to ensure a good fit to data over a high level of sparsity. For a detailed discussion on the impact and interpretation of *δ*_ABC_, we refer the reader to Joergensen et al., 2023 ((Jørgensen et al., 2023)).

In the SLInG runs discussed in the results section, we thus set *δ*_abc_ = 0.0125. In addition to the value of *δ*_ABC_ set by the user, SLInG includes a threshold for the largest possible value that the distance measure *d* might take for the initial guess of the parameter values. SLInG will autonomously readjust *δ*_ABC_ based on this threshold to facilitate convergence. However, after the initial burn-in phase, SLInG will re-evaluate the choice of *δ*_ABC_ and set it to the value chosen by the user if this value is consistent with the threshold for the best-fitting model within the burn-in. Here, we set the threshold to 35, i.e. SLInG will use *δ*_abc_ = 0.0125 if the distance measure of the best-fitting model within the burn-in divided by 0.0125 is lower than 35.

To evaluate the impact of the choice for *δ*_ABC_, we repeated the analysis for a subset of ODE models using different values of *δ*_ABC_. We operated three SLInG runs for the two antipodal models, the one with all ORs (1) and the one with all ANDs (20), for four additional values of *δ*_ABC_, obtained by multiplying the original value of *δ*_ABC_ (0.0125) by 0.5, 2, 4, and 8. The SLInG algorithm itself automatically increases the value of *δ*_ABC_ if the precision for the *δ*_ABC_ set cannot be achieved, which happens in the first three runs for model 1 and in the second run for model 20 (Figure S4). We computed the error distance *d* setting the initial values of *δ*_ABC_ to *{*0.00625, 0.0125, 0.025, 0.05, 0.1*}* (Figure S4).

We obtained very similar errors and links when dividing or multiplying *δ*_abc_ by a factor of 2. When setting *δ*_abc_ = 0.00625, *δ*_abc_ = 0.0125 and *δ*_abc_ = 0.025, the final errors are thus very similar within each model, indicating that the algorithm is quite stable in a neighbourhood of the value we picked (*δ*_abc_ = 0.0125). However, when dividing by 2, we note that this result is partly due to the fact that the low value of *δ*_abc_ comes into conflict with the aforementioned threshold. Indeed, when setting *δ*_abc_ = 0.00625 in model 1, SLInG resets the value to 0.01 in all three SLInG runs. For model 20, on the other hand, the value is only slightly increased to 0.0075 once.

When looking at higher values of *δ*_abc_ (0.05 and 0.1), the errors start to increase significantly, while the number of non-zero parameters decreases. This behaviour is to be expected: By increasing *δ*_abc_, SLInG puts less weight on the data and thus strives towards a sparser fit, i.e. the algorithm discards more parameters at the expense of larger errors (see also Jørgensen et al., 2023). For the five different values of *δ*_abc_ explored in Figure 4C, the average number of non-zero parameters is 7.2, 6.8, 6.8, 3.8, and 3.2, respectively, listed in the order of increasing *δ*_abc_. Note that when only 3 parameters are kept in any SLInG run, these three parameters are identifiable as the basal productions of the three genes, i.e. no network is present. Hence, for *δ*_abc_ = 0.1, we indeed obtained a very sparse solution in which no links are present.

## Acknowledgements

AB is supported by the BBSRC. AC is supported by the EPSRC. ACSJ is supported by the Eric and Wendy Schmidt AI in Science Postdoctoral Fellowship, a Schmidt Futures program.

## Author Contributions

AB and AC conducted the majority of the experimental and computational work. MH and ACSJ contributed key data, code and analysis. VS and MB supervised the work. AB, AC, ACSJ, MH, MB, and VS wrote and edited the manuscript.

**Supplementary Figure 1:**
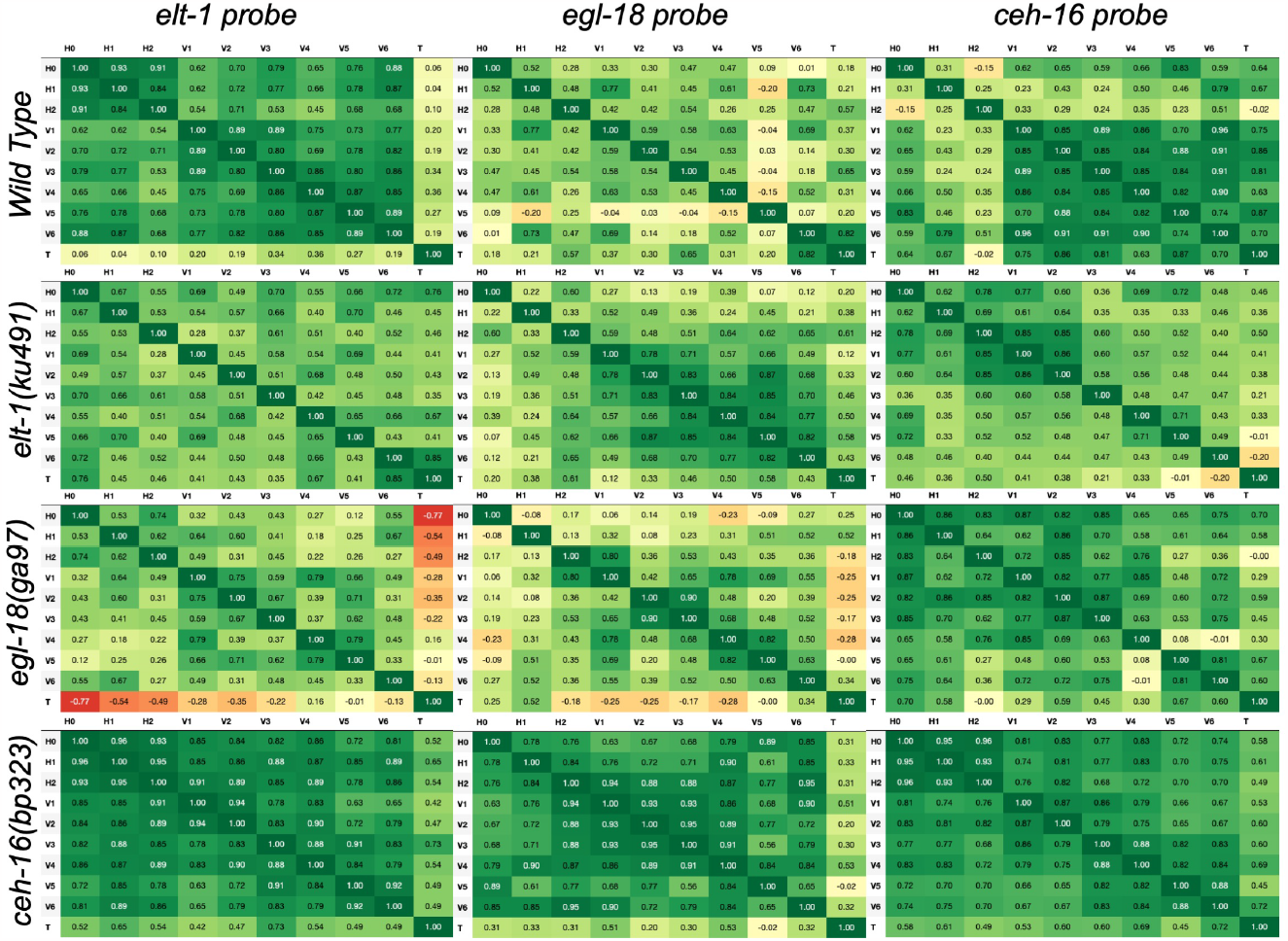
Correlation matrices of smFISH data of the core seam cell genes between all the seam cells under wildtype and mutant conditions: These matrices show the correlation of the expression of each core gene, in each cell, under mutant and wildtype conditions, to the same gene in every other cell in the same animal. Green denotes a strong positive correlation, and red denotes a strong negative correlation

**Supplementary Figure 2:**
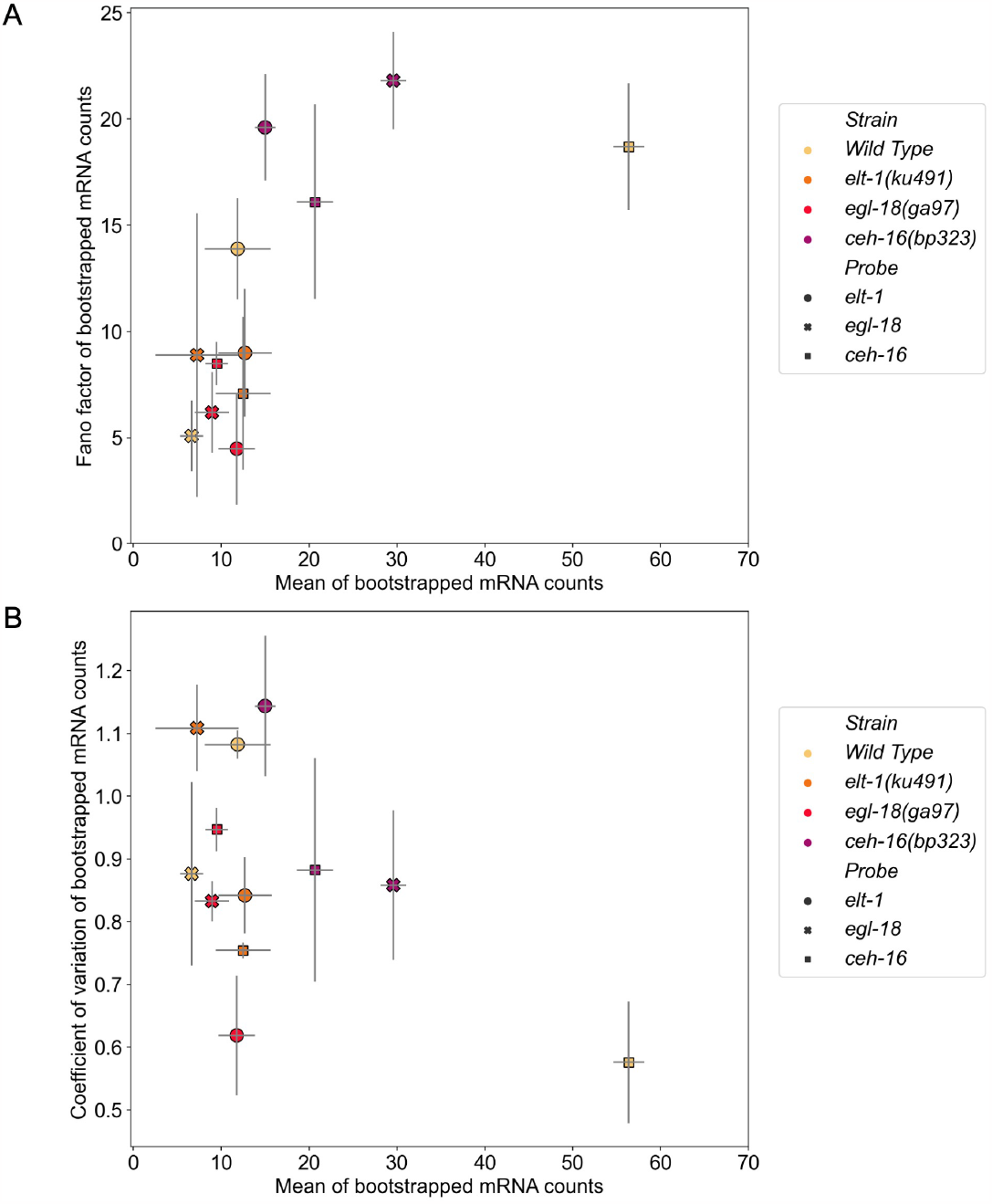
Noise in smFISH data of the core genes under wildtype and mutant conditions. **A:** The Fano factors (calculated as 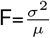) of smFISH data quantifying the variability of the core seam cell gene expression relative to the mean of each bootstrapping case. The error bars here illustrate all bootstrapping cases that fall within the inter-quartile range of the data. **B:** The coefficient of variation (calculated as 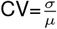) of the bootstrapped smFISH data showing how the standard deviation of the data changes with respect to the mean. The error bars indicate the interquartile range of all the bootstrapped data.

**Supplementary Figure 3:**
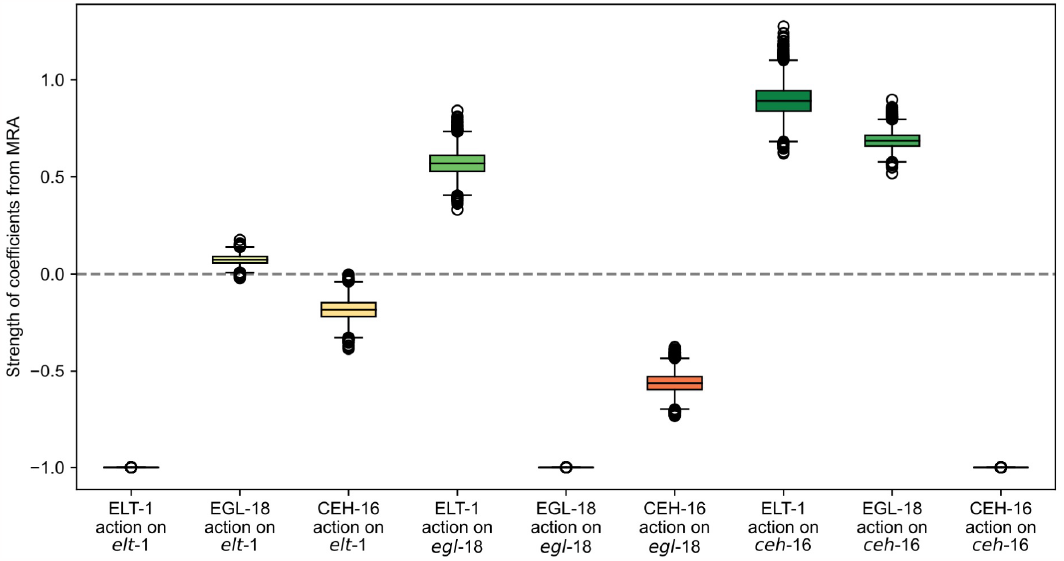
Interaction matrices resulting from the Modular Response Analysis. The matrix that is generated when all cells are used as opposed to V1-V4. The element in position (i, J) (row, COLUMN) indicates the effect of protein J on gene i. Positive numbers correspond to activation, negative to repression, and values act as an indicator of the strength of the action.

**Supplementary Figure 4:**
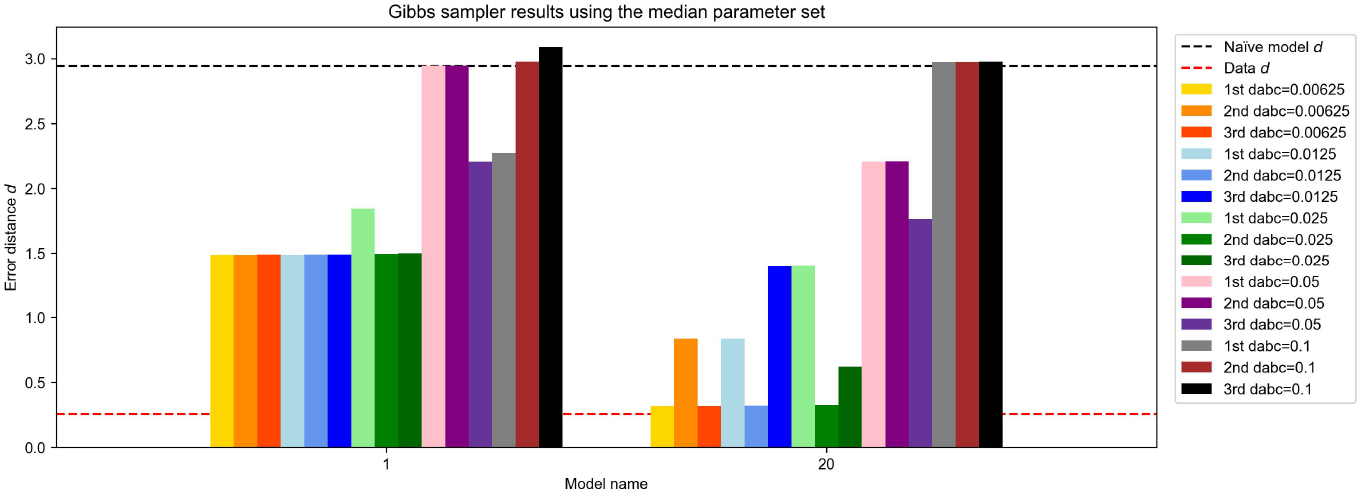
Testing the models with different *δ*_ABC_. We compare the behaviour of the two antipodal models (1 with all OR links and 20 with all AND links) under different *δ*_ABC_.

**Supplementary Table 1:**
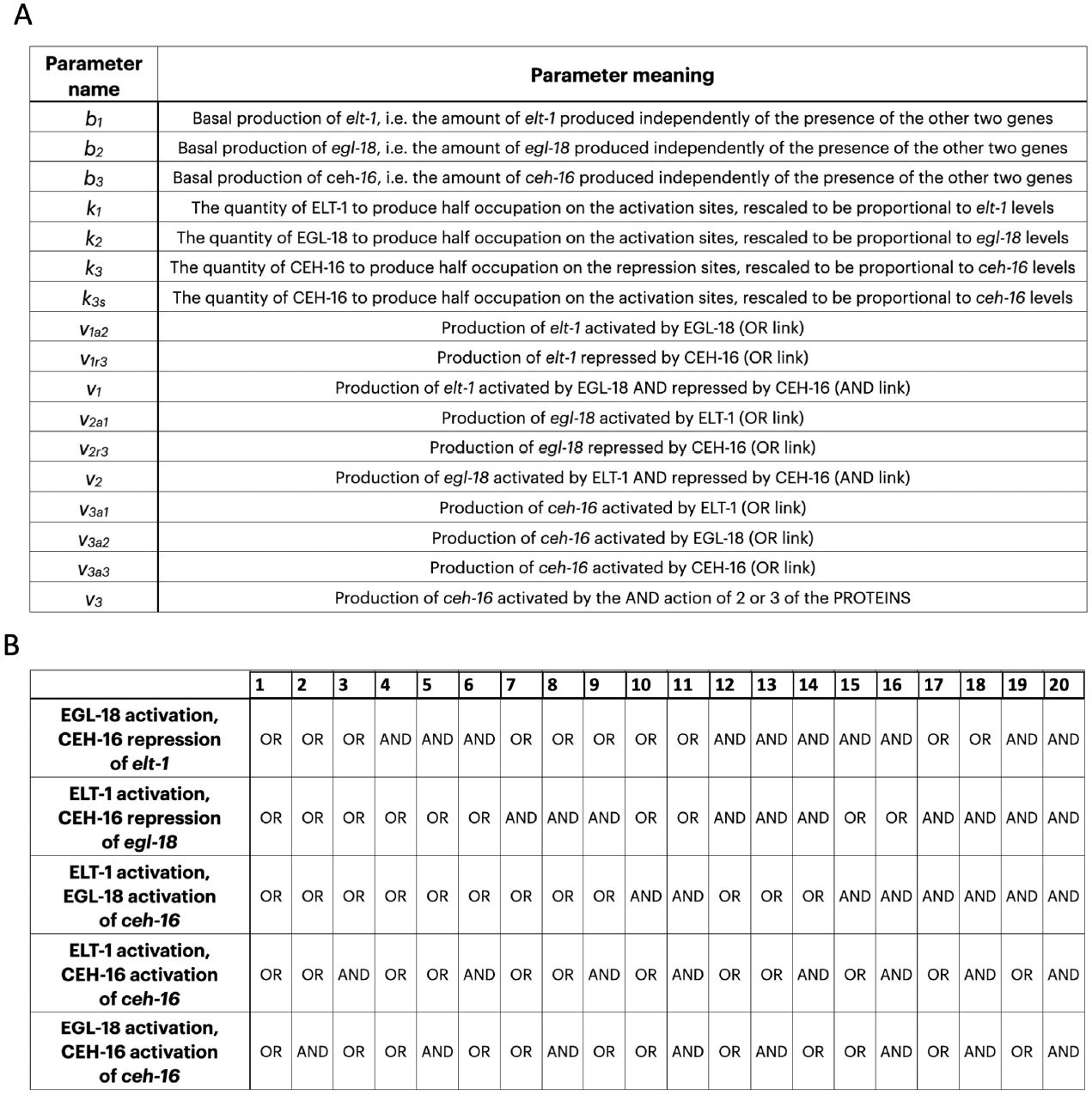
Additional description of the various ODE models used. **A:** Parameter names and definition. **B:** Description of the AND/OR logic of different models tested based on the initial MRA.

**Supplementary Table 2:**
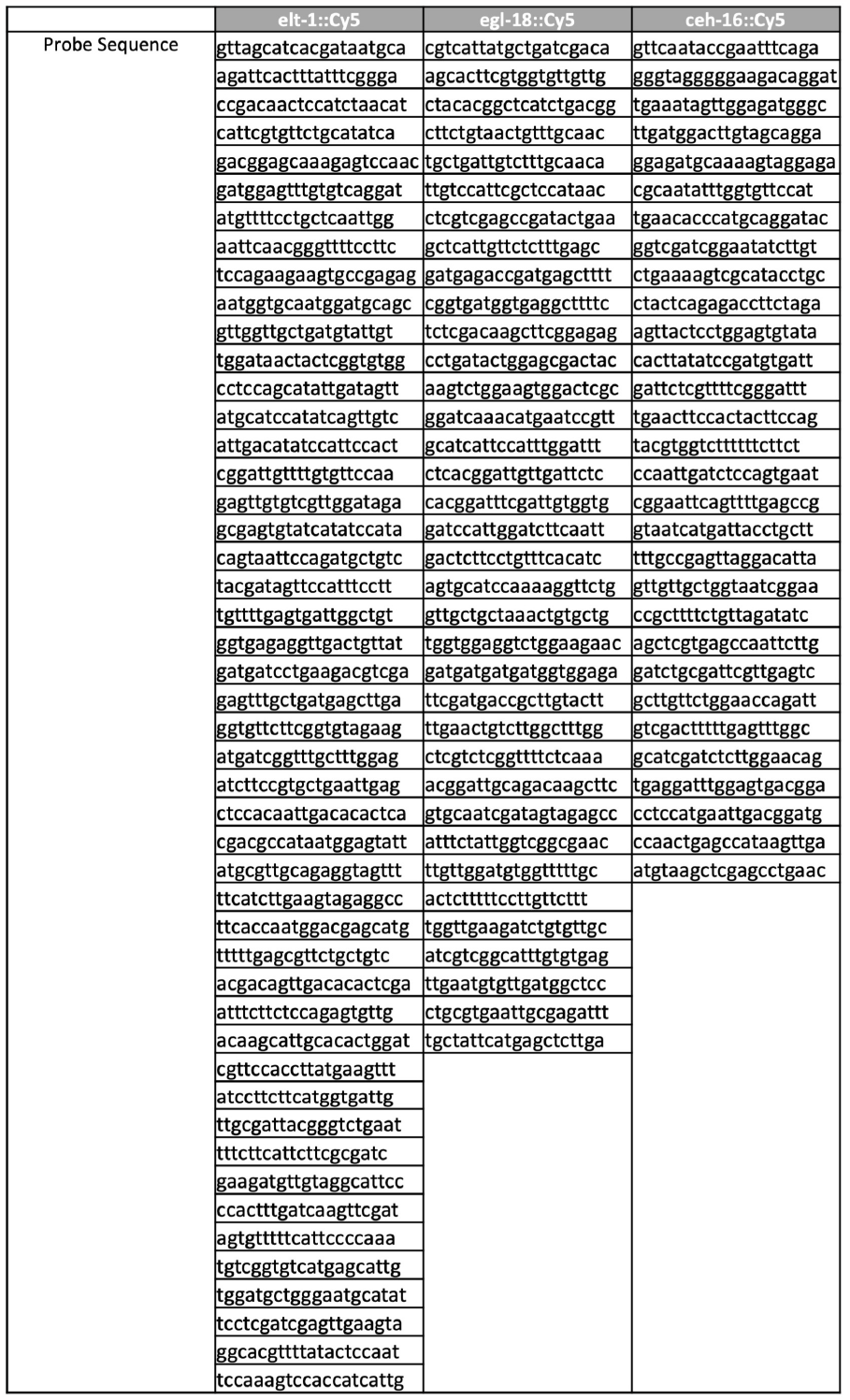
smFISH probes used in this study: The full sequence of the cy5 bound smFISH probes used to quantify *elt-1, egl-18* and *ceh-16*.

## References

1. A. A. Alahmadi, J. A. Flegg, D. G. Cochrane, C. C. Drovandi, and J. M. Keith. A comparison of approximate versus exact techniques for bayesian parameter inference in nonlinear ordinary differential equation models. Royal Society Open Science, 7(3), 2020. ISSN 20545703. doi: 10.1098/rsos.191315.

2. C. Alexandre and J.-P. Vincent. Requirements for transcriptional repression and activation by engrailed in drosophila embryos. Development, 130(4):729–739, 2003. pmid:12506003.

3. G. Balázsi, A. V. Oudenaarden, and J. J. Collins. Cellular decision making and biological noise: from microbes to mammals. Cell, 144 (6):910–925, 2011. pmid:21414483.

4. M. Barkoulas, J. S. van Zon, J. Milloz, A. van Oudenaarden, and M.-A. Félix. Robustness and epistasis in the C. elegans vulval signaling network revealed by pathway dosage modulation. Developmental Cell, 24(1):64–75, 2013.

5. L. R. Baugh, J. C. Wen, A. A. Hill, D. K. Slonim, E. L. Brown, and C. P. Hunter. Synthetic lethal analysis of caenorhabditis elegans posterior embryonic patterning genes identifies conserved genetic interactions. Genome biology, 6(5):1–8, 2005.

6. M. A. Beaumont. Approximate bayesian computation. Annual review of statistics and its application, 6:379–403, 2019.

7. G. J. Both, S. Choudhury, P. Sens, and R. Kusters. Deepmod: Deep learning for model discovery in noisy data. Journal of Computational Physics, 428(1), 2021. ISSN 10902716. doi: 10.1016/j.jcp.2020.109985.

8. C. Brabin, P. J. Appleford, and A. Woollard. The Caenorhabditis elegans gata factor elt-1 works through the cell proliferation regulator bro-1 and the fusogen eff-1 to maintain the seam stem-like fate. PLoS genetics, 7(8), 2011. pmid:21829390.

9. S. Brenner. The genetics of Caenorhabditis elegans. Genetics, 77(1):71–94, 1974. pmid:4366476.

10. S. L. Brunton, J. L. Proctor, and J. N. Kutz. Discovering governing equations from data by sparse identification of nonlinear dynamical systems. Proceedings of the national academy of sciences, 113(15):3932–3937, 2016.

11. G. Cassata, G. Shemer, P. Morandi, R. Donhauser, B. Podbilewicz, and R. Baumeister. ceh-16/engrailed patterns the embryonic epidermis of Caenorhabditis elegans. Development, 132(4):739–749, 2005. pmid:15659483.

12. X. Chen, Z. Jane Wang, and M. J. McKeown. A Bayesian Lasso via reversible-jump MCMC. Signal Processing, 91(8):1920–1932, 2011. ISSN 01651684. doi: 10.1016/j.sigpro.2011.02.014.

13. M. L. Cohen, S. Kim, K. Morita, S. H. Kim, and M. Han. The gata factor elt-1 regulates C. elegans developmental timing by promoting expression of the let-7 family micrornas. PLoS Genet, 11(3):e1005099, 2015.

14. E. P. Consortium. An integrated encyclopedia of dna elements in the human genome. Nature, 489(7414):57, 2012. pmid:22955616.

15. V. Davies, R. Reeve, W. T. Harvey, F. F. Maree, and D. Husmeier. A sparse hierarchical bayesian model for detecting relevant antigenic sites in virus evolution. Computational Statistics, 32(3):803–843, 2017. ISSN 16139658. doi: 10.1007/s00180-017-0730-6.

16. M. J. Dunlop, R. S. C. III, J. H. Levine, R. M. Murray, and M. B. Elowitz. Regulatory activity revealed by dynamic correlations in gene expression noise. Nature genetics, 40(12):1493–1498, 2008.

17. D. M. Eisenmann and S. K. Kim. Protruding vulva mutants identify novel loci and wnt signaling factors that function during Caenorhabditis elegans vulva development. Genetics, 156(3):1097–1116, 2000. pmid:11063687.

18. M. B. Elowitz, A. J. Levine, E. D. Siggia, and P. S. Swain. Stochastic gene expression in a single cell. Science, 297(5584):1183–1186, 2002.

19. T. S. Gardner and J. J. Faith. Reverse-engineering transcription control networks. Physics of life reviews, 2(1):65–88, 2005.

20. M. Genestine, L. Lin, M. Durens, Y. Yan, Y. Jiang, S. Prem, K. Bailoor, B. Kelly, P. K. Sonsalla, and P. G. Matteson. Engrailed-2 (en2) deletion produces multiple neurodevelopmental defects in monoamine systems, forebrain structures and neurogenesis and behavior. Human molecular genetics, 24(20):5805–5827, 2015.

21. L. Gorrepati, K. W. Thompson, and D. M. Eisenmann. C. elegans gata factors egl-18 and elt-6 function downstream of wnt signaling to maintain the progenitor fate during larval asymmetric divisions of the seam cells. Development, 140(10):2093–2102, 2013. pmid:23633508.

22. M. Hecker, S. Lambeck, S. Toepfer, E. V. Someren, and R. Guthke. Gene regulatory network inference: data integration in dynamic models—a review. Biosystems, 96(1):86–103, 2009.

23. M. Hintze, S. L. Koneru, S. P. Gilbert, D. Katsanos, J. Lambert, and M. Barkoulas. A cell fate switch in the caenorhabditis elegans seam cell lineage occurs through modulation of the wnt asymmetry pathway in response to temperature increase. Genetics, 214(4): 927–939, 2020.

24. X. Huang, E. Tian, Y. Xu, and H. Zhang. The C. elegans engrailed homolog ceh-16 regulates the self-renewal expansion division of stem cell-like seam cells. Developmental biology, 333(2):337–347, 2009.

25. A. C. S. Jørgensen, M. Sturrock, A. Ghosh, and V. Shahrezaei. Bayesian model discovery for reverse-engineering biochemical networks from data. bioRxiv, 2023. doi: 10.1101/2023.09.15.557764.

26. P. M. Joshi, M. R. Riddle, N. J. Djabrayan, and J. H. Rothman. Caenorhabditis elegans as a model for stem cell biology. Developmental dynamics: an official publication of the American Association of Anatomists, 239(5):1539–1554, 2010.

27. G. Karlebach and R. Shamir. Modelling and analysis of gene regulatory networks. Nature reviews Molecular cell biology, 9(10): 770–780, 2008.

28. D. Katsanos, S. L. Koneru, L. M. Boukhibar, N. Gritti, R. Ghose, P. J. Appleford, M. Doitsidou, A. Woollard, J. S. van Zon, and R. J. Poole. Stochastic loss and gain of symmetric divisions in the c. elegans epidermis perturbs robustness of stem cell number. Plos Biology, 15(11):e2002429, 2017.

29. B. N. Kholodenko, A. Kiyatkin, F. J. Bruggeman, E. Sontag, H. V. Westerhoff, and J. B. Hoek. Untangling the wires: a strategy to trace functional interactions in signaling and gene networks. Proceedings of the National Academy of Sciences, 99(20):12841–12846, 2002. pmid:12242336.

30. E. Klipp, R. Herwig, A. Kowald, C. Wierling, and H. Lehrach. Systems biology in practice: concepts, implementation and application. John Wiley & Sons, 2005.

31. K. Koh and J. H. Rothman. Elt-5 and elt-6 are required continuously to regulate epidermal seam cell differentiation and cell fusion in C. elegans. Development, 128(15):2867–2880, 2001. pmid:11532911.

32. K. Koh, S. M. Peyrot, C. G. Wood, J. A. Wagmaister, M. F. Maduro, D. M. Eisenmann, and J. H. Rothman. Cell fates and fusion in the C. elegans vulval primordium are regulated by the egl-18 and elt-6 gata factors are apparent direct targets of the lin-39 hox protein. Development, 129(22):5171–5180, 2002. pmid:12399309.

33. S. L. Koneru. Investigating the role of the fusogen eff-1 and natural genetic variation in caenorhabditis elegans seam cell development. Imperial College London, 2020.

34. S. L. Koneru, F. X. Quah, R. Ghose, M. Hintze, N. Gritti, J. S. van Zon, and M. Barkoulas. A role for the fusogen eff-1 in epidermal stem cell number robustness in caenorhabditis elegans. Scientific reports, 11(1):1–13, 2021.

35. S. Kramer, E. Meyer-Natus, C. Stigloher, H. Thoma, A. Schnaufer, and M. Engstler. Parallel monitoring of rna abundance, localization and compactness with correlative single molecule fish on lr white embedded samples. Nucleic acids research, 49(3):e14, 2021.

36. M. M. Kudron, A. Victorsen, L. Gevirtzman, L. W. Hillier, W. W. Fisher, D. Vafeados, M. Kirkey, A. S. Hammonds, J. Gersch, and H. Ammouri. The modern resource: genome-wide binding profiles for hundreds of drosophila and caenorhabditis elegans transcription factors. Genetics, 208(3):937–949, 2018.

37. Y. Luo, B. C. Hitz, I. Gabdank, J. A. Hilton, M. S. Kagda, B. Lam, Z. Myers, P. Sud, J. Jou, and K. Lin. New developments on the encyclopedia of dna elements (encode) data portal. Nucleic acids research, 48(D1):D882–D889, 2020.

38. H. Mallick and N. Yi. A new bayesian lasso. Statistics and its Interface, 7(4):571–582, 2014. ISSN 19387997. doi: 10.4310/SII.2014.v7.n4.a12.

39. H. Markel, J. Chandler, and W. Werr. Translational fusions with the engrailed repressor domain efficiently convert plant transcription factors into dominant-negative functions. Nucleic acids research, 30(21):4709–4719, 2002.

40. D. Mercatelli, L. Scalambra, L. Triboli, F. Ray, and F. M. Giorgi. Gene regulatory network inference resources: A practical overview. Biochimica et Biophysica Acta (BBA)-Gene Regulatory Mechanisms, 1863(6):194430, 2020.

41. N. Moris, C. Pina, and A. M. Arias. Transition states and cell fate decisions in epigenetic landscapes. Nature Reviews Genetics, 17 (11):693–703, 2016.

42. B. Munsky, B. Trinh, and M. Khammash. Listening to the noise: random fluctuations reveal gene network parameters. Molecular systems biology, 5(1):318, 2009.

43. D. L. Nelson, A. L. Lehninger, and M. M. Cox. Lehninger principles of biochemistry. Macmillan, 2008.

44. B. D. Page, W. Zhang, K. Steward, T. Blumenthal, and J. R. Priess. Elt-1, a gata-like transcription factor, is required for epidermal cell fates in Caenorhabditis elegans embryos. Genes & development, 11(13):1651–1661, 1997. pmid:9224715.

45. T. Park and G. Casella. The bayesian lasso. Journal of the American Statistical Association, 103(482):681–686, 2008. ISSN 01621459. doi: 10.1198/016214508000000337.

46. K. Petratou, T. Subkhankulova, J. A. Lister, A. Rocco, H. Schwetlick, and R. N. Kelsh. A systems biology approach uncovers the core gene regulatory network governing iridophore fate choice from the neural crest. PLoS genetics, 14(10):e1007402, 2018.

47. A. Raj, P. V. D. Bogaard, S. A. Rifkin, A. V. Oudenaarden, and S. Tyagi. Imaging individual mrna molecules using multiple singly labeled probes. Nature methods, 5(10):877–879, 2008.

48. V. Shahrezaei and P. S. Swain. The stochastic nature of biochemical networks. Current opinion in biotechnology, 19(4):369–374, 2008.

49. J. A. Smith, P. McGarr, and J. S. Gilleard. The caenorhabditis elegans gata factor elt-1 is essential for differentiation and maintenance of hypodermal seam cells and for normal locomotion. Journal of cell science, 118(24):5709–5719, 2005.

50. J. Spieth, Y. H. Shim, K. Lea, R. Conrad, and T. Blumenthal. elt-1, an embryonically expressed Caenorhabditis elegans gene homologous to the gata transcription factor family. Molecular and cellular biology, 11(9):4651–4659, 1991. pmid:1875944.

51. M. P. Stumpf. Inferring better gene regulation networks from single-cell data. Current Opinion in Systems Biology, 27:100342, 2021.

52. J. E. Sulston and H. R. Horvitz. Post-embryonic cell lineages of the nematode, Caenorhabditis elegans. Developmental biology, 56 (1):110–156, 1977.

53. M. Sáez, R. Blassberg, E. Camacho-Aguilar, E. D. Siggia, D. A. Rand, and J. Briscoe. Statistically derived geometrical landscapes capture principles of decision-making dynamics during cell fate transitions. Cell Systems, 13(1):12–28. e3, 2022. pmid:34536382.

54. K. W. Thompson, P. Joshi, J. S. Dymond, L. Gorrepati, H. E. Smith, M. W. Krause, and D. M. Eisenmann. The paired-box protein pax-3 regulates the choice between lateral and ventral epidermal cell fates in C. elegans. Developmental biology, 412(2):191–207, 2016.

55. B. M. Turner and T. Van Zandt. Hierarchical approximate bayesian computation. Psychometrika, 79(2):185–209, 2014. ISSN 00333123. doi: 10.1007/s11336-013-9381-x.

