## Supplementary figures and images for "Inference of the core gene regulatory network underlying seam cell development in *Caenorhabditis elegans*"

### Supplementary Figure 1

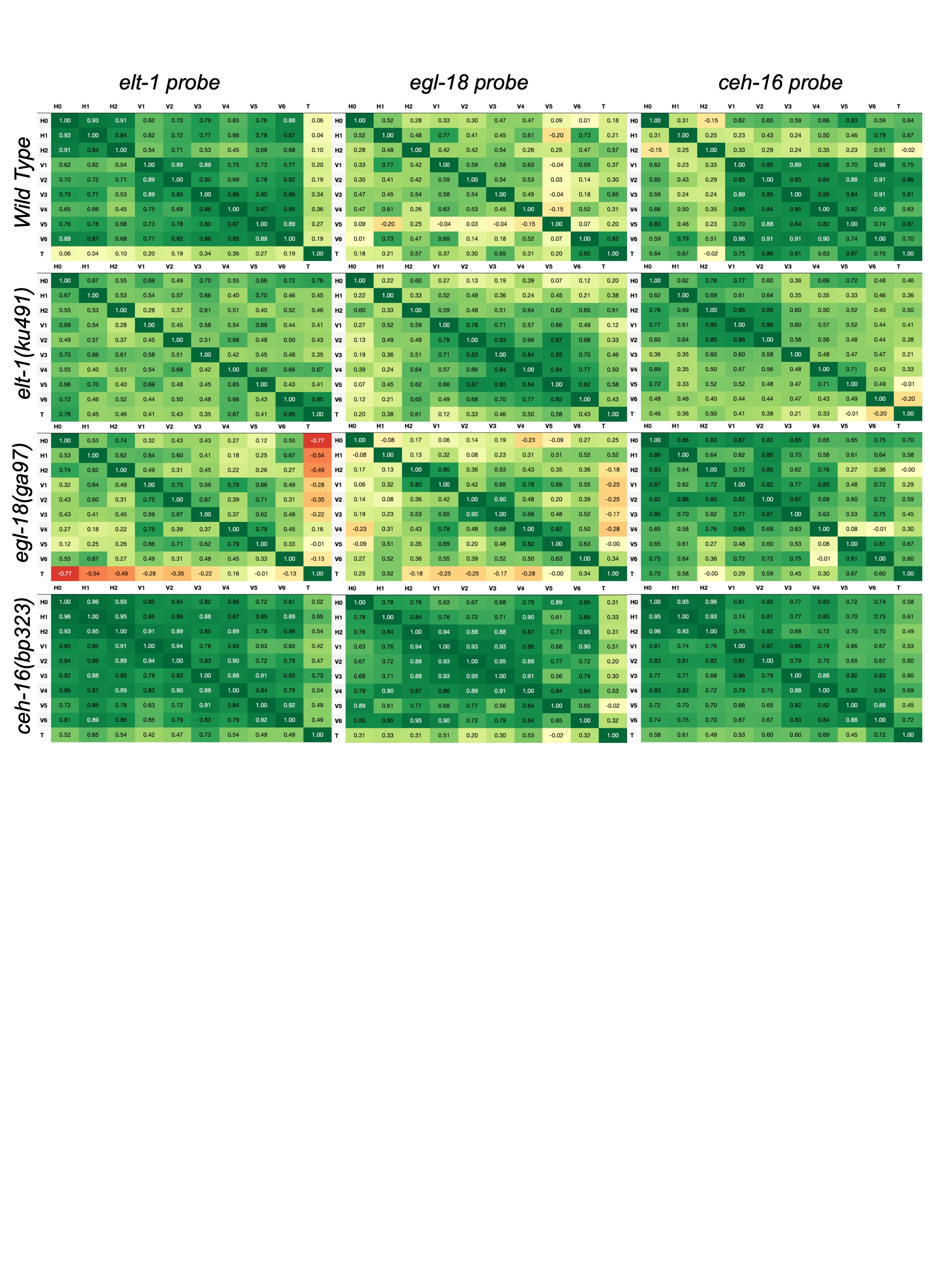

### Supplementary Figure 2

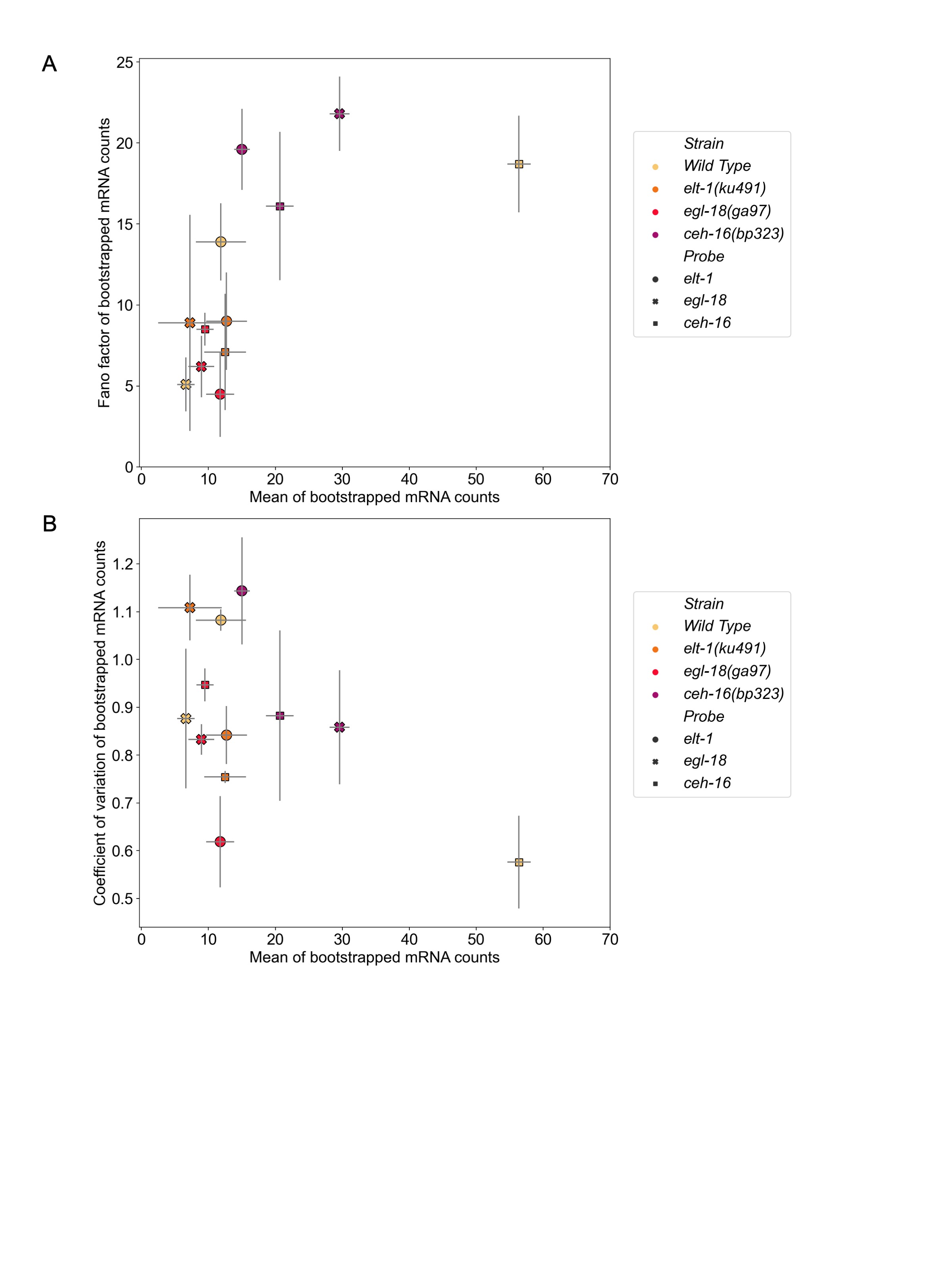

### Supplementary Figure 3

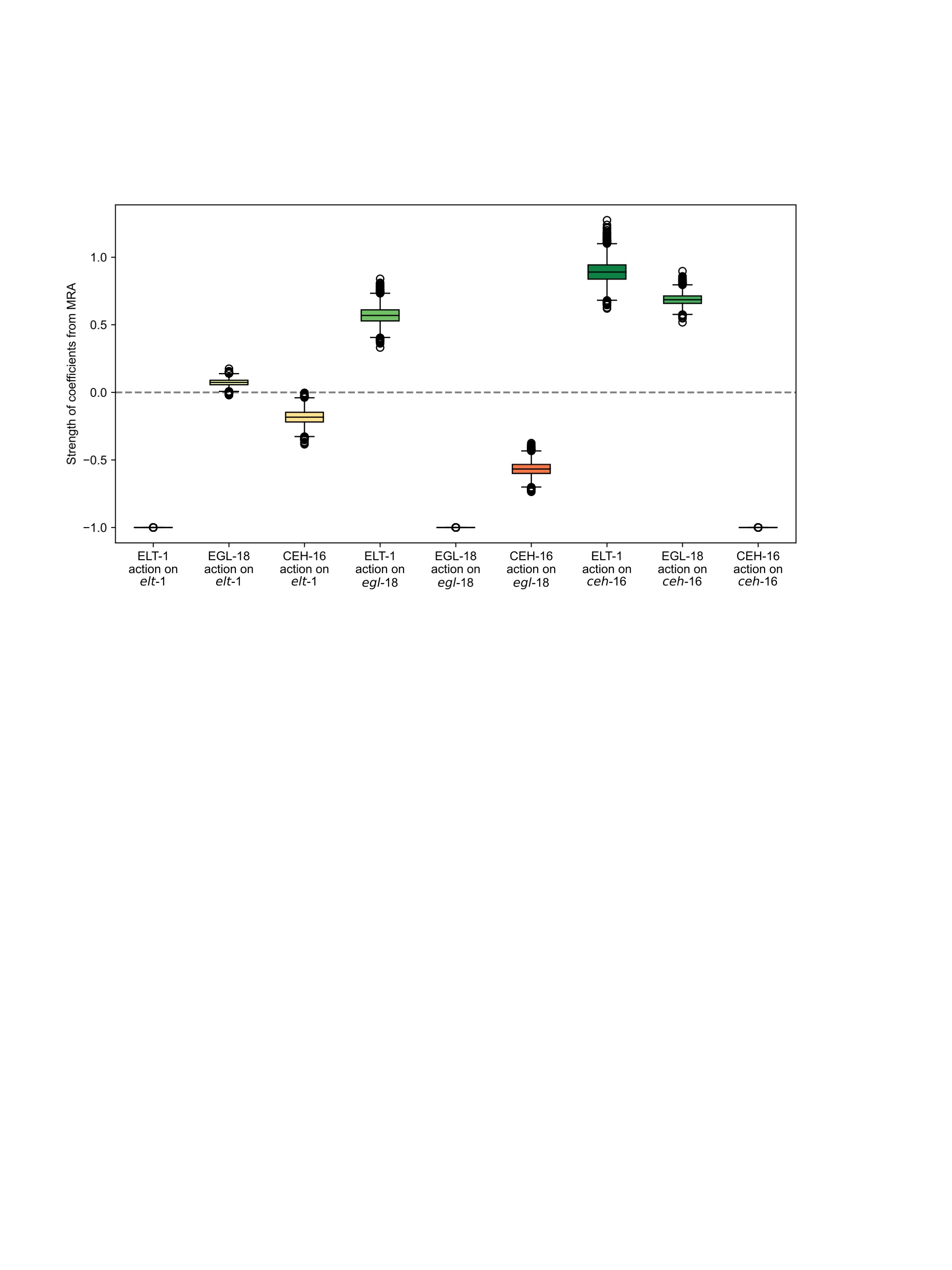

### Supplementary Figure 4

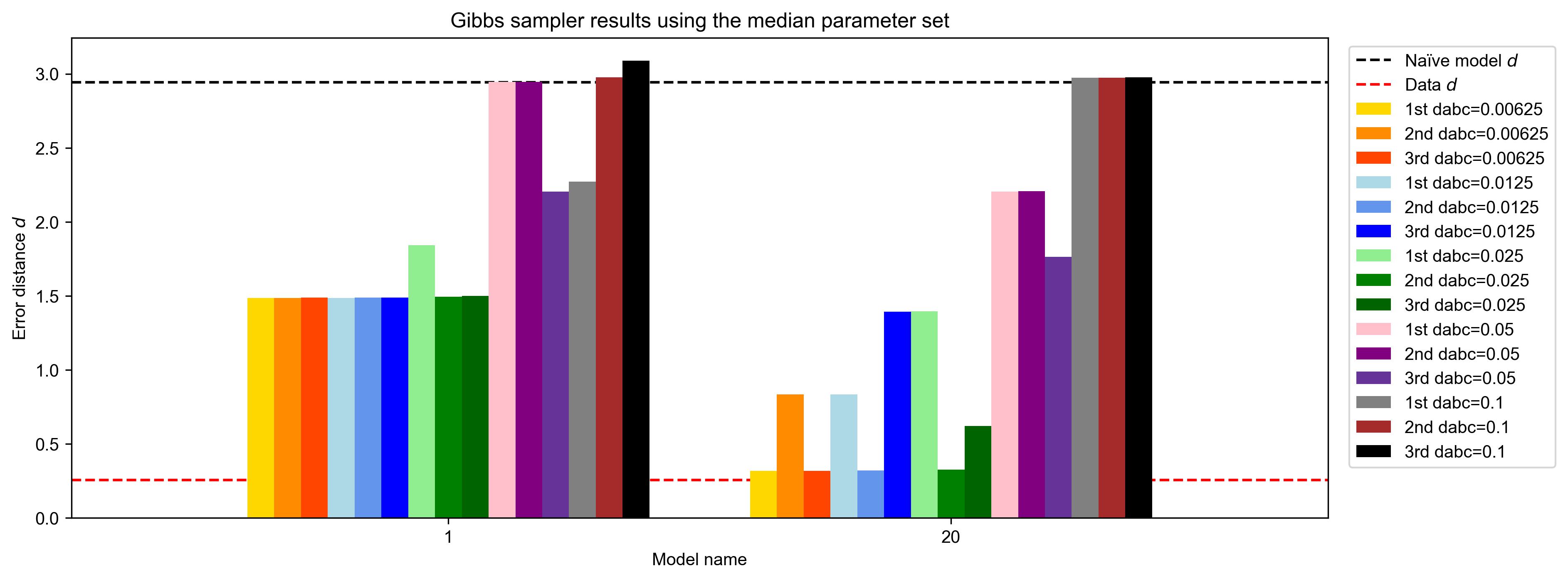

### Supplementary Table 1

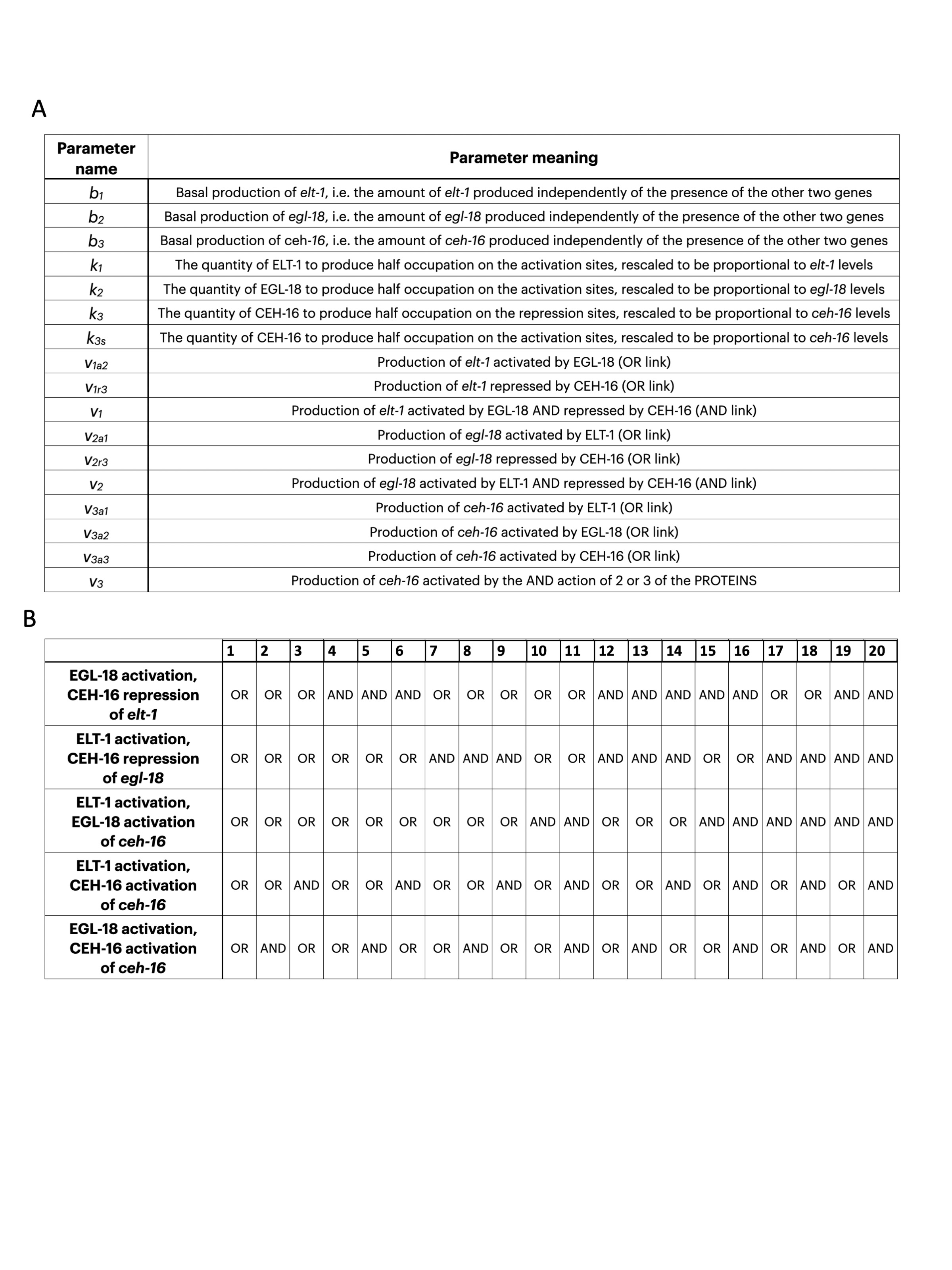

### Supplementary Table 2

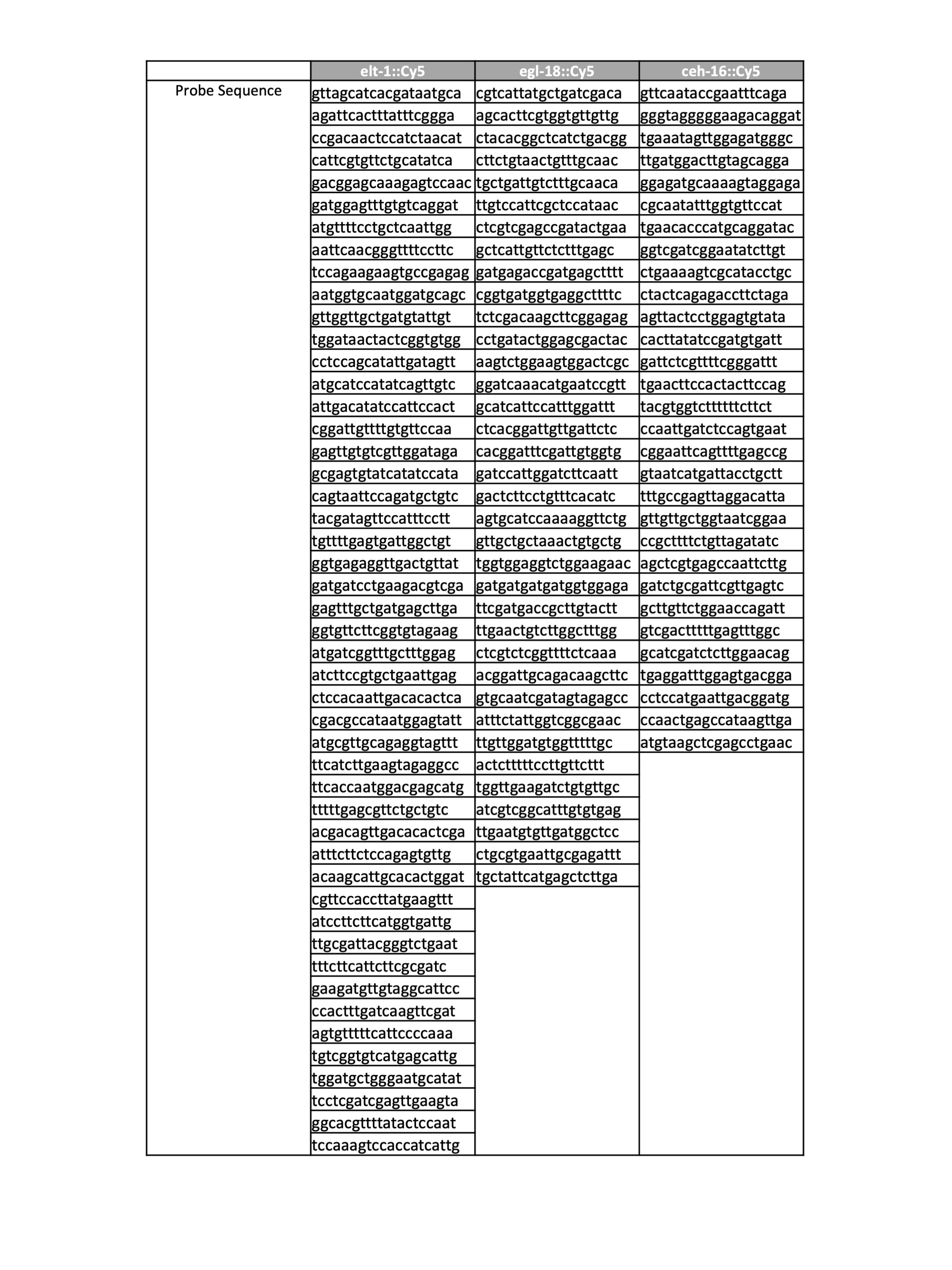
